# Model-guided engineering of DNA sequences with predictable site-specific recombination rates

**DOI:** 10.1101/2021.08.02.454698

**Authors:** Qiuge Zhang, Samira M. Azarin, Casim A. Sarkar

**Affiliations:** Department of Chemical Engineering and Materials Science, University of Minnesota, Minneapolis, Minnesota 55455, United States; Department of Biomedical Engineering, University of Minnesota, Minneapolis, Minnesota 55455, United States

**Keywords:** site-specific DNA recombination, reaction rate prediction, high-throughput selection, machine learning

## Abstract

Site-specific recombination (SSR) is an important tool in genome editing and gene circuit design. However, its applications are limited by the inability to simply and predictably tune SSR reaction rates across orders of magnitude. Facile rate manipulation can in principle be achieved by modifying the nucleotide sequence of the DNA substrate of the recombinase, but the design principles for rationally doing so have not been elucidated. To enable predictable tuning of SSR reaction kinetics via DNA sequence, we developed an integrated experimental and computational method to parse individual nucleotide contributions to the overall reaction rate, which we used to analyze and engineer the DNA attachment sequence attP for the inversion reaction mediated by the serine recombinase Bxb1. A quantitative PCR method was developed to measure the Bxb1 reaction rate *in vitro*. Then, attP sequence libraries were designed, selected, and sequenced to inform a machine-learning model, which revealed that the Bxb1 reaction rate can be accurately represented assuming independent contributions of nucleotides at key positions. Next, we used the model to predict the performance of DNA site variants in reaction rate assays both *in vitro* and in *Escherichia coli*, with flipping rates ranging from 0.01- to 10-fold that of the wild-type attP sequence. Finally, we demonstrate that attP variants with predictable DNA recombination rates can be used in concert to achieve kinetic control in gene circuit design by coordinating the coexpression of two proteins in both their relative proportion and their total amount. Our high-throughput, data-driven method for rationally tuning SSR reaction rates through DNA sequence modification enhances our understanding of recombinase function and expands the synthetic biology toolbox.

**Graphical abstract:** 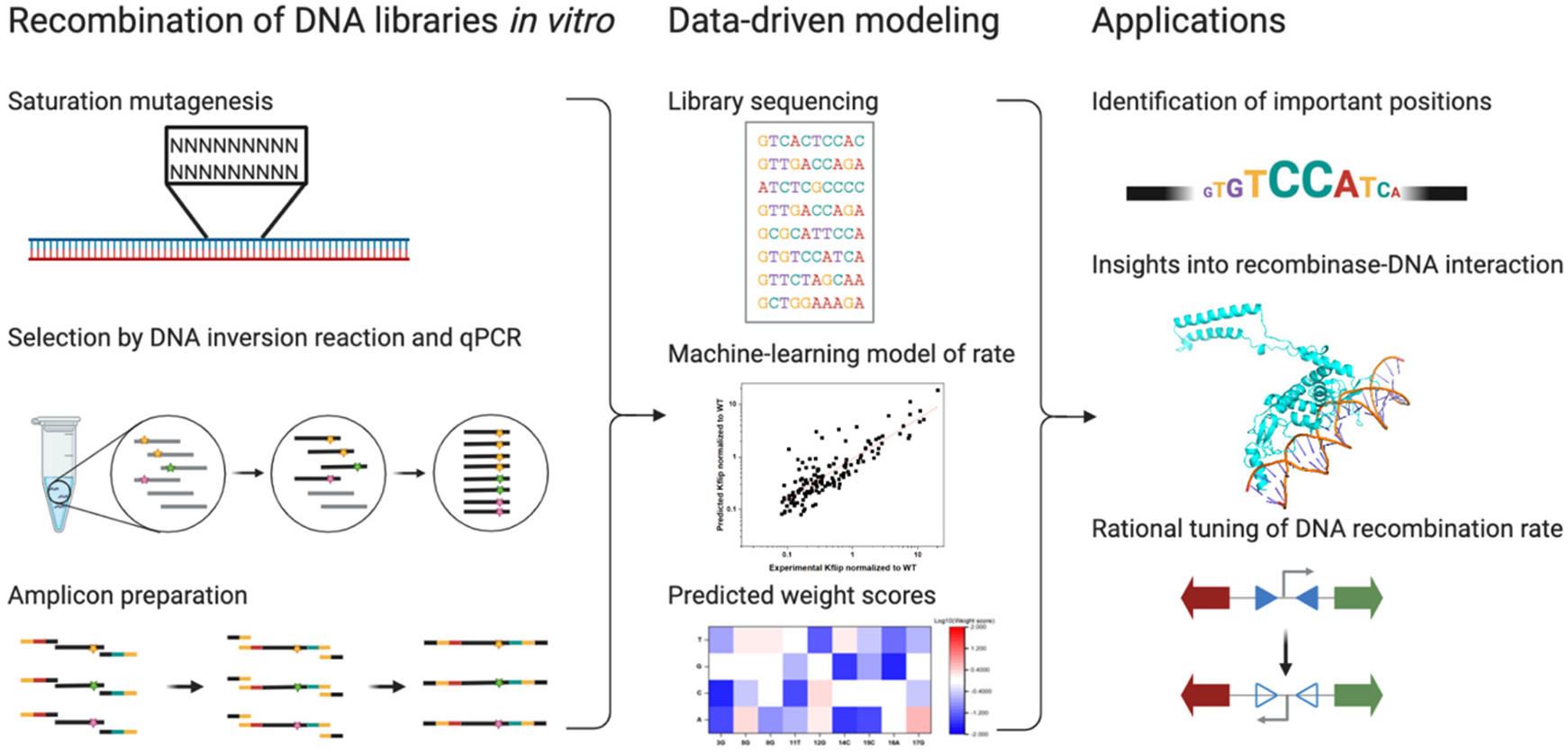

## Introduction

Site-specific recombination (SSR) technology relies on recombinases that can precisely recognize two DNA sites, form an intermediate complex, cut, swap, and recombine the DNA in a new configuration, resulting in gene insertions, deletions, or inversions.^1^ Based on the active residue in the catalytic domain, the recombinase superfamily is divided into two groups: tyrosine recombinases and serine recombinases.^2^ Each group can be further subdivided by directionality (bidirectional/unidirectional) for tyrosine recombinases or by size (large/small) for serine recombinases.^3^ Among these four recombinase subgroups, the large serine recombinases (LSRs) are considered one of the most powerful tools in synthetic biology based on the following properties:^4^

- Irreversibility: LSRs have non-identical recognition sites typically known as attB (attachment site bacteria) and attP (attachment site phage) and yield hybrid product sites attL and attR. LSRs cannot target the hybrid attL and attR sites to regenerate attP and attB, resulting in an exceptionally stable DNA recombination product, which is in contrast to the commonly used Cre-lox and FLP-FRT systems.^5^ This feature is important in applications such as human cell genome editing^6^ and gene circuits for data storage in living cells.^7^
- Simplicity: In contrast to some tyrosine recombinases such as λ integrase, which requires long attP sites (>200 bp), supercoiled DNA structure, and other factors to stabilize DNA bending, serine recombinases have short DNA sites (<50 bp) and no required DNA topology or cofactors.^8^
- Efficiency: LSRs such as Bxb1, TP901 and PhiC31 have been demonstrated to be efficient in both prokaryotic and eukaryotic cells, in gene therapy,^9^ memory circuit design,^10,11^ and genome editing.^12^

However, the lack of fine control over SSR reaction rates has hindered the use of this technology in regulating DNA recombination in gene circuits. An understanding of the DNA determinants of such processes could not only lead to improvements in wild-type recombination rates but could also provide a suite of parts that could be coupled together to enable higher-order information processing in genetic circuits via kinetic control. Here, we focused on understanding and engineering the DNA inversion reaction mediated by the mycobacteriophage integrase Bxb1^13^, though given the shared functional mechanisms of LSRs, our approach should be readily translatable to other LSRs as well.

Previous engineering approaches for regulating biochemical reaction rates has focused on altering key amino acid residues in the enzyme.^14^ However, rational protein design is limited by the lack of high-resolution recombinase-DNA complex crystal structures. To our knowledge, only one structure of a DNA-bound LSR, Listeria phage A118 integrase, has been reported and the resolution of the protein-DNA interface is not sufficiently high.^15^ Despite ongoing efforts to understand the interactions between amino acid residues of recombinases and nucleotides, static recombinase-DNA complex crystal structures cannot provide sufficient information to understand DNA sequence determinants of SSR rates.^15–19^ In addition to direct protein-DNA contacts, charge/shape complementarity and water-mediated interactions contribute to the SSR rate of different DNA sequences.^20^ Furthermore, mutations in the recombinase can alter protein stability or solubility, confounding efforts to rationally tune reaction rates via enzyme engineering. Therefore, we instead focused on engineering the DNA attachment sites and developing a method to rationally design DNA attachment site sequences to modulate the SSR reaction rate. Previously, the impact of single and double base substitutions in the Bxb1 attP site on specificity and directionality was reported.^21^ Another group used a high-throughput approach to identify DNA specificity determinants by selecting saturated mutagenized DNA site libraries.^22^ Although essential for better understanding the recombination mechanism and revealing potential off-target substrates, these studies focused only on DNA sequence specificity. Considering the potential application of LSRs in the design of genetic circuits, it would also be useful to be able to rationally tune their reaction kinetics. Through this new mode of kinetic control at the DNA level, it would be possible to use recombinases in applications beyond genetic memory, such as coordination of protein expression dynamics and temporal ordering of genetically encoded processes.

We therefore sought to develop a method to programmably tune the Bxb1 reaction rate via the DNA attachment sequences. However, a method to accurately measure SSR reaction rates or a platform to screen DNA sequences for recombination on a large scale has not been well established. In this study, we developed a qPCR-based method for quantifying relative SSR reaction rate as well as a platform for profiling SSR reaction rates of selected sequences from a designed DNA library using next-generation sequencing (NGS). Then, we constructed a data-driven model to quantify the contributions of different nucleotide substitutions in attP to the overall SSR reaction rate. Finally, using assays both *in vitro* and in *E. coli*, we demonstrated accurate model predictions of rates of DNA inversion. Our study enables rational modulation of SSR reaction rates, providing a new form of kinetic control for predictably tuning synthetic genetic circuits and gene therapies.

## Results and Discussion

### DNA library design

For our initial DNA library design, we first identified nucleotide positions in the attP and attB DNA attachment sites of Bxb1 that could potentially be substituted to vary the reaction rate while maintaining recognition specificity. As shown in Fig. 1A, during the SSR reaction, the attP and attB attachment sites each bind a Bxb1 dimer. After synapsis through the interaction between the coiled-coil (CC) motifs at the different sites, the DNA sequence is cleaved at the center and then rotated 180°. After rotation, the conformation of the CC groups bound together on the same DNA strand is much more thermodynamically stable; thus, the ligation step is essentially irreversible and drives the overall reaction from the attP and attB substrate sites to the attL and attR product sites.^23^ Notably, the attP and attB sites for Bxb1 have non-identical sequences, with both having two quasi-half-sites attP-L/attP-R and attB-L/attB-R. A Bxb1 monomer bound to the attP-L halfsite has direct contact with the DNA site sequence via the zinc ribbon domain (ZD), the recombinase domain (RD), and the linker connecting ZD and RD (Fig. 1B).^15^ The shorter length of the attB site forces the linker to adopt a different conformation when bound to attB half sites.^15^ As shown in Fig. 1B (right panel, underlined), the four half-sites attP-L, attP-R, attB-L, and attB-R have ~50% conserved bases. Previous studies have demonstrated that the homology positions were highly conserved for specificity.^24^ Therefore, we hypothesized that these conserved positions may directly interact with protein residues and are necessary for DNA recognition, whereas bases at other asymmetric positions might be substituted to vary reaction rates.

**Figure 1.**
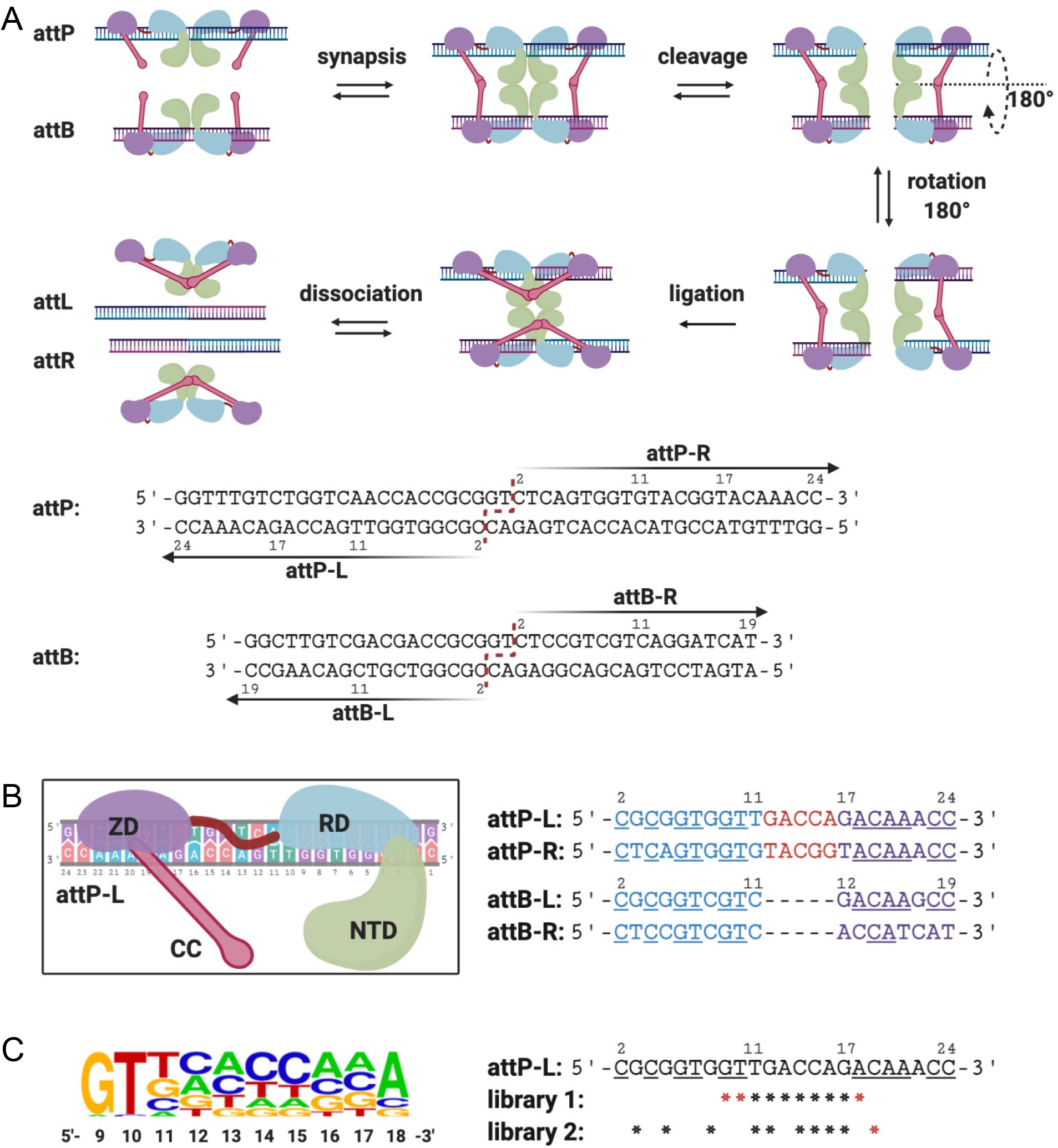
Bxb1 attP-L DNA library design. (A) Top: DNA inversion mediated by Bxb1 recombinase. Attachment sites attP and attB are bound by Bxb1 dimers, followed by synapsis through intermolecular binding of CC motifs. DNA sequences are then cleaved in the middle by the recombinase domain and rotated 180°. The resulting sticky ends, which are complementary, are ligated. After rotation, Bxb1 forms a more stable conformation, with CC motifs interacting within the same Bxb1 dimer on a given DNA sequence. Finally, Bxb1 dissociates from the two newly generated attachment sites, attL and attR, and the intervening DNA sequence is inverted. Bottom: DNA sequences of Bxb1 attP and attB sites. Cleavage happens at the central bases GT in both (red dashed line). Both attP and attB have two quasi-palindromic half sites attP-L/attP-R and attB-L/attB-R (also referred to as attP/attP’and attB/attB’ half-sites). (B) Left: one Bxb1 monomer includes the N-terminal domain (NTD, green), recombinase domain (RD, blue), zinc ribbon domain (ZD, purple), and coiled-coil domain (CC, pink). A linker (red) connects RD and ZD and adopts different conformations when bound to attP or attB half-sites due to their differential lengths. Right: comparison of attP and attB sites and their features. Four half-site sequences are aligned from center to end, with numbering from the 5’-terminus to the 3’-terminus. Underlined nucleotides indicate positions that are conserved across at least three half sites. Positions bound to RD or ZD are shown in blue or purple, respectively, and bases in red are aligned with the linker. (C) Left: Weblogo of base frequency from position 9 to position 18 in library 1 after an *E. coli*-based selection. Right: designed saturation mutagenesis libraries 1 and 2, where an asterisk represents a degenerate position in the library (with equal representation of bases A, C, G, and T at each such position; Supplemental Method 1 and Table S4).

To test this hypothesis, we chose to perform saturation mutagenesis on the attP-L half-site based on the following considerations. First, the attP site has more asymmetric positions than attB (nucleotides in red, Fig. 1B), potentially expanding the tunable range of reaction rates. Second, considering limitations of transformation efficiency and NGS read depth in the selection step, the number of positions for saturation mutagenesis had to be limited to 10 nucleotides (library size = 4^10^ ~ 1 million). Lastly, within the attP site, substituting bases in both the attP-L and attP-R half-sites would be more likely to result in a loss of specificity.^21^ We therefore opted to keep the attP-R half-site unchanged, introducing mutations only within the attP-L half-site.

In previous studies, base conservation at homologous positions was characterized *in vitro* by gel electrophoresis.^22^ Since SSR can be impacted by multiple factors, including direct protein-DNA contacts and long-distance interactions, we designed a selection method in *E. coli* to ensure that our selected attP-L variants function *in vivo*. This selection method entails inducing a DNA inversion reaction that confers expression of chloramphenicol acetyltransferase, with selected attP-L variants then recovered by plating on chloramphenicol-containing agar plates (Fig. S1). We constructed a DNA library containing random bases at 10 continuous positions from 9 to 18 in attP-L, including symmetric sites and asymmetric sites (library 1 in Fig. 1B). From initial selections with library 1 in *E. coli*, 20 colonies with SSR functionality were sequenced and three highly conserved bases – 9G, 10T, and 18A – were identified (Fig. S1 and Supplemental Method 1). These conserved bases are shared by at least three out of the four wild-type half-sites (Fig. 1B), suggesting that these positions are essential for maintaining SSR specificity, while other positions 11T, 12G, 13A, 14C, 15C, 16A, 17G were tolerant to nucleotide substitutions.

Given these observations, we hypothesized that the homologous positions among the four half-sites are indispensable for SSR recognition, while the asymmetric positions could provide more flexibility in tuning the reaction rates. We therefore designed a new saturation mutagenesis DNA library (library 2, Fig. 1C) to identify variants with a broader range of reaction rates by altering the asymmetric positions (3G, 5G, 8G, 11T, 12G, 14C, 15C, 16A, 17G), while keeping one putatively conserved base, 19C, as a control in the selection.

### DNA flipping rate assay and library selection in vitro

Traditional methods for characterizing DNA recombination events are based on differential electrophoretic mobility of substrates and products,^18,21,23^ but they are not amenable to high-throughput experiments. Furthermore, quantification of band intensities in gels is not sensitive enough to detect low product concentrations at the beginning of the reaction, when the initial rate can capture differences in the intrinsic efficiency of DNA variants. In this work, we developed a qPCR-based method to accurately measure the initial SSR reaction rate by quantifying the recombinant DNA product concentration. As shown in Fig. 2A, two primers were designed to selectively amplify the flipped product DNA but not the substrate. Briefly, we constructed a linear DNA fragment flanked by attP and attB sites positioned in opposite directions. Bxb1 binds to attP/attB sites and then inverts the flanked DNA segment. Our designed primers are complementary to the adjacent sequences of the cleavage site on the attP sequence, and the inversion reaction changes both the orientation of one of the primers and the DNA strand to which it anneals, such that PCR results in exponential amplification of the inverted template, which, importantly, includes the attP-L sequence. For unflipped DNA substrate, both primers extend in the same direction on the same strand, so no amplification is possible by PCR. We used qPCR to measure the percentage of flipping with different Bxb1 concentrations for wild-type (WT) attP sites, and the result was consistent with those from DNA gel electrophoresis (Fig. S2). To test the sensitivity of this quantification method, a standard curve of Cq values and flipped DNA template concentration was plotted (Fig. 2B). The total copy number of unflipped and flipped DNA fragments was fixed at 10^9^ per qPCR reaction volume (20 μl) and the ratio of flipped to unflipped DNA was varied (0:1, 1:10^7^, 1:10^6^, 1:10^5^, 1:10^4^, 1:10^3^, 1:10^2^, 1:10, 1:0). As shown in Fig. 2B, our qPCR approach accurately measured the flipped DNA percentage (r^2^ = 0.998). Additionally, the lower bound of the linear range was ~10^3^ flipped DNA copies, indicating that this method has a high sensitivity and can detect a flipping percentage as low as 0.0001%. Compared with previous methods based on differences in electrophoretic mobility^21^ or exonuclease digestion,^22^ our method is facile and can accurately measure initial reaction rates.

**Figure 2.**
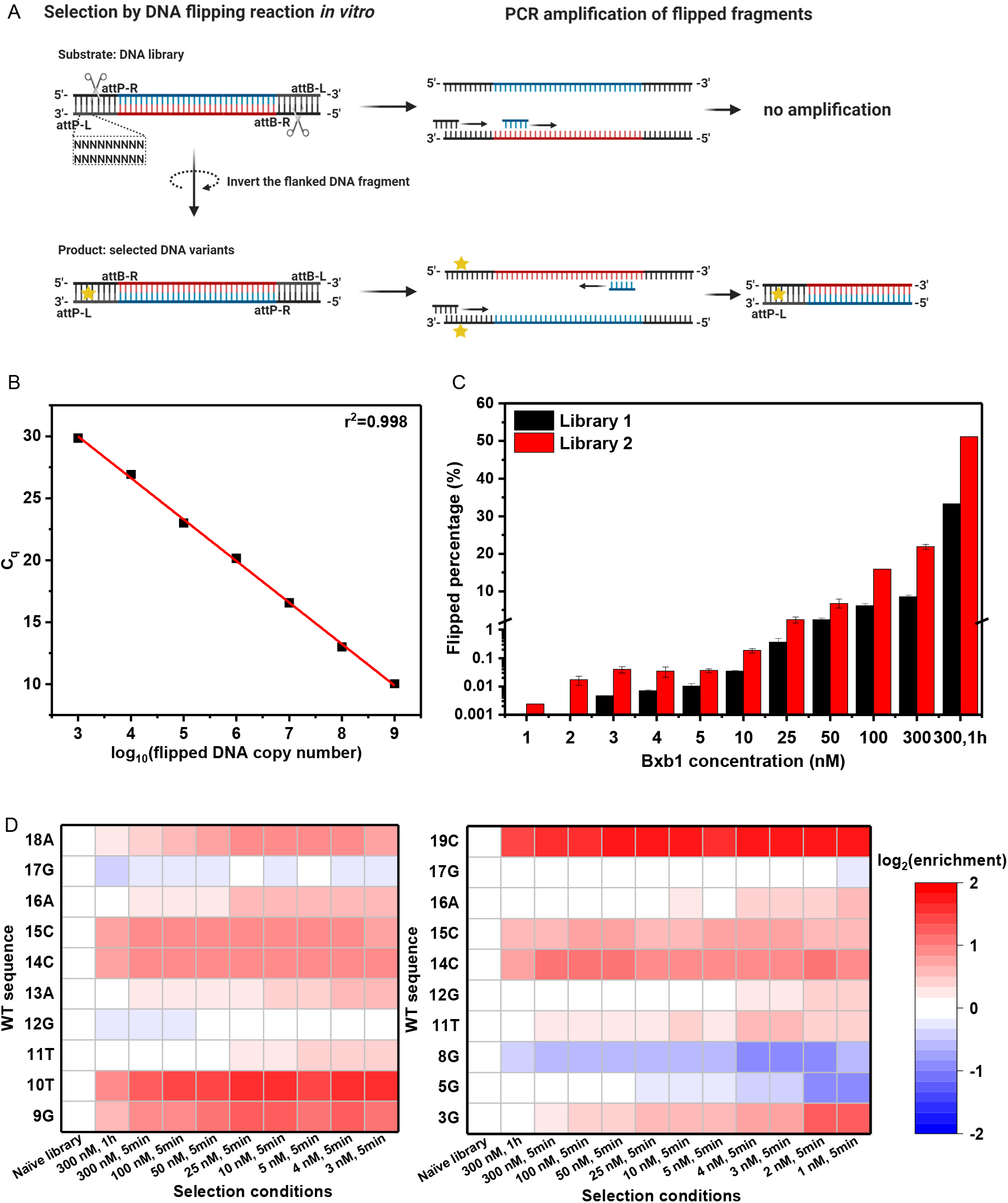
Quantification of DNA inversion and attP-L library selections by qPCR. (A) Library selection procedure. First, a DNA site-saturation library is constructed by designed oligonucleotides. Second, sequences with DNA recombination functionality (yellow star) undergo inversion in the reaction solution. Last, the flipped sequences are selectively enriched by qPCR. For flipped DNA fragments, the black and blue primers anneal to two different DNA strands, resulting in amplified products (bottom); for unflipped DNA, the primers anneal to the same strand and no PCR product can be produced (top). (B) Standard curve for qPCR quantification of flipped DNA fragments. The templates for qPCR are mixtures of flipped and unflipped DNA with different ratios and the same total copy number 10^9^ (slope = −3.378, intercept = 40.169, r^2^=0.998). (C) qPCR quantification of DNA library flipping. Bxb1 at concentrations varying from 1 nM to 300 nM was added to the reaction buffer with 10 nM DNA library and then incubated at 30 °C for 5 min. The reaction was terminated by denaturation at 80 °C for 20 min. The sample with 300 nM Bxb1 and incubated for 1 h at 30 °C was considered the reaction with the most permissive conditions. (D) Heat map of wild-type nucleotide evolution as a function of selection stress, which increases from left to right in each map (left: library 1; right: library 2).

With this sensitive and accurate qPCR-based rate assay, we selected for DNA variants with high recombination efficiencies using a DNA library as the substrate. To prepare the library, we used oligonucleotides containing a degenerate base (N = equimolar A, C, G, and T) at designated positions (Fig. 2A). Then, recombinant Bxb1 was added to the DNA library and incubated in reaction buffer. The attP-L variants with efficient SSR functionality underwent flipping of the DNA sequence flanked by the attP and attB sites, whereas the remaining inefficient or nonfunctional DNA molecules stayed unflipped. From this mixture, the flipped DNA product was selectively amplified by qPCR and sequenced by NGS. Sequence variants that led to faster reaction rates produced more flipped DNA products and thus appeared more frequently in the NGS pool. By ranking the frequency of the flipped DNA sequences, we obtain a list of sequences in order of their flipping rate.

To optimize the selection conditions, we tested Bxb1 concentrations ranging from 1 nM to 300 nM. At high enzyme concentrations, we were unable to distinguish the reaction rates of attP-L variants due to the lower selection pressure. Further, low enzyme concentrations resulted in very low flipped percentages, even below the sensitivity of qPCR detection. As shown in Fig. 2C, qPCR was unable to detect flipped DNA in library 1 at the lowest Bxb1 concentrations (1 nM and 2 nM), whereas under the same conditions, 0.002% and 0.017% of flipped DNA substrate was detected in library 2. The flipped percentages for library 2 were higher than library 1 at all other Bxb1 concentrations as well. This observation was consistent with our hypothesis in library design, as library 2 (with only one putatively conserved position, 19C) was expected to have more functional sequences than library 1 (with three conserved positions, 9G, 10T, and 18A). Last, in measuring the flipped percentage under the most permissive reaction conditions (300 nM Bxb1 and 1 h incubation time), we found that 33% of library 1 and 51% of library 2 variants possessed SSR competence. Although Bxb1 is considered to have high specificity, this result suggests that the half-site attP-L is not highly resistant to nucleotide substitution when paired with three unchanged WT half-sites, attP-R, attB-L, and attB-R. More importantly, it revealed the potential to modulate DNA recombination rate by altering the bases at specific positions in attP-L.

After sequencing the flipped libraries using NGS, we quantified the percentage of wild-type bases at each nucleotide position (Table S5 and S6) and plotted them as an enrichment heat map. The enrichment score for a given base was defined as the ratio between its percentage in the selected library and its percentage in the naïve library. As shown in Fig. 2D, the high enrichment scores of WT bases at positions 9, 10, and 19 supported our hypothesis that homologous positions tended to be functionally conserved. However, for other positions, especially 5, 8, 12, and 17, WT bases were more amenable to substitution, with some substitutions even preferred over WT bases under harsh selection conditions (Tables S5 and S6). As shown in Fig. 2D, these results demonstrated that our *in vitro* selection approach using qPCR and NGS was able to identify optimal base substitutions, consistent with our preliminary *in vivo* selections in *E. coli*. In addition, these substitutions were more pronounced at lower Bxb1 concentrations. Next, to investigate the relationship between the entire attP-L sequence and its corresponding SSR reaction rate, we analyzed the frequency of individual attP-L variants from the NGS results and then converted the frequency into an SSR rate constant, k_fl_ip.

### Model training and calculation of weight scores

For each individual attP variant, we counted the frequency of occurrence from the refined NGS results. As shown in Fig. 3A, as the Bxb1 concentration decreased, fewer variants were selected and appeared in the sequencing results, and their frequencies varied more between the different selected DNA sequences due to the same depth of 5×10^5^ reads during sequencing. To demonstrate that frequency and reaction rate have the correlation we expected, we selected 8 sequences, including WT, from the 3000 most frequently occurring sequences and performed SSR assays on each individual sequence. For these tested sequences, the frequencies in the NGS results and the flip percentages in the SSR assays showed a good correlation, r^2^ = 0.83 (Fig. S3C). Given this experimentally confirmed relationship, we converted the occurrence frequency to a more biologically relevant value, flipping rate constant (k_flip_), to represent the intrinsic activity of the attP-L variants (Supplementary Methods 2). In brief, under selection conditions of low Bxb1 and DNA concentrations, SSR reactions with different DNA variants as substrates can be considered independent, with the reaction rate for an individual sequence governed by a first-order reaction with a rate constant k_flip_. Thus, for each individual attP-L variant, its frequency of occurrence within the total reads was proportional to its percentage of flipped DNA after a period of reaction time, so its flipping rate can be calculated from its k_flip_ value and a reaction time of 5 min.

**Figure 3.**
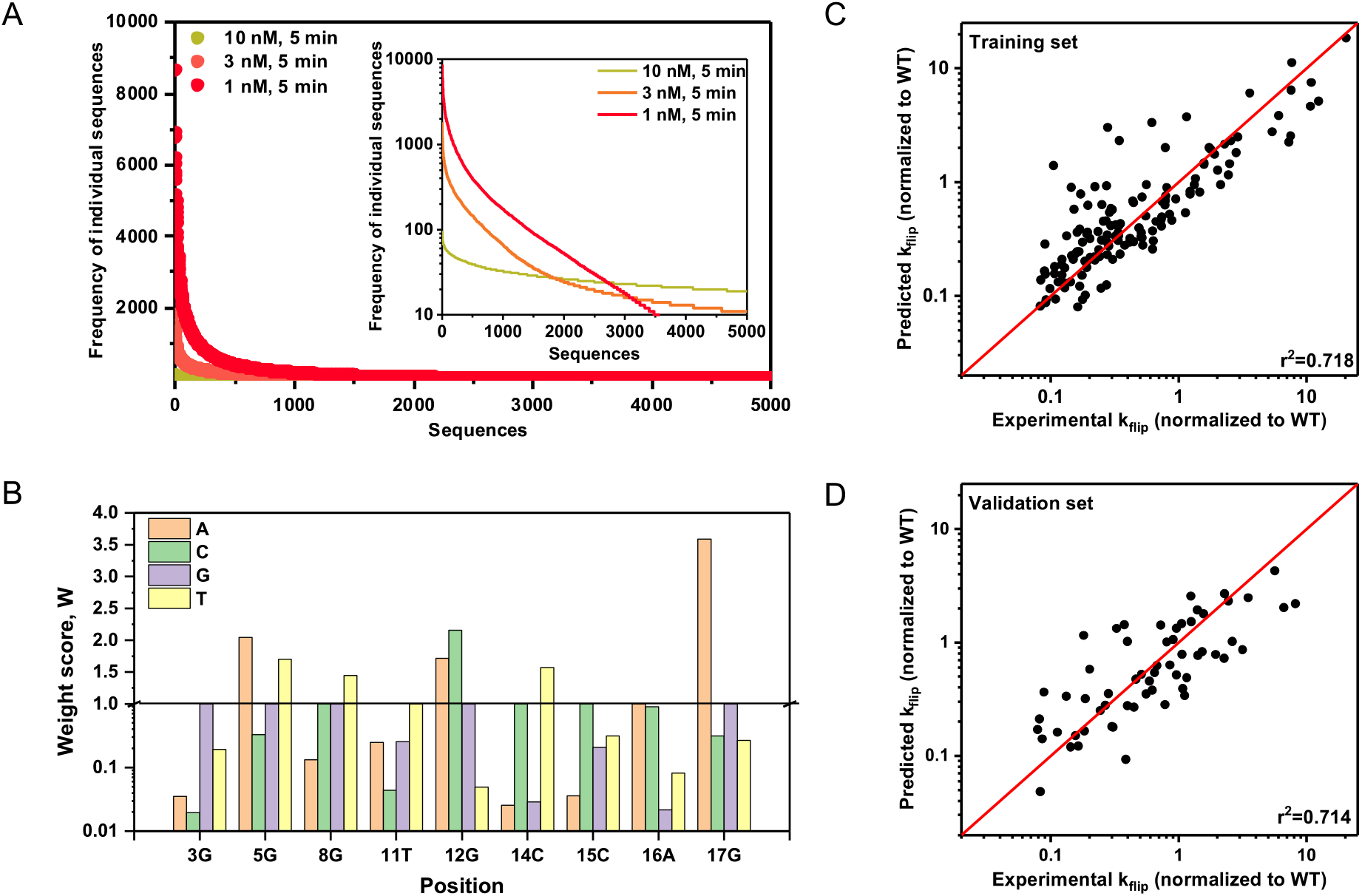
Model fitting of NGS result. (A) Occurrence frequency of individual sequences selected with different Bxb1 concentrations. (B) Weight scores generated from linear regression. Experimental log (k_flip_) values are derived from the occurrence frequency in NGS results (Supplementary Methods 2). Weight scores of the reference WT sequences are fixed as 1 (i.e., W_i1_=1). For the y-axis in 3B, 0.01-1 is on a log_10_ scale and 1-4 is on a linear scale. Predicted k_flip_ values from the model correlate with the experimental k_flip_ values for (C) 140 randomly chosen training sequences (70% of 200 sequences) and (D) 60 randomly chosen validation sequences (30% of 200 sequences).

Interestingly, the approximately linearly decreasing trend when frequency curves were displayed on a logarithmic scale (Fig. 3A) implied that the logarithm of the k_flip_ of a sequence might be reasonably predicted by a model that assumes independent contributions of each DNA base in the attP-L sequence. To test our hypothesis and quantify the contributions of different bases at each nucleotide position, we built a linear model and defined weight score parameters to quantify the relative contribution of each possible nucleotide substitution. In this model, the input data were the DNA sequences and the output data were the sequence-specific rate constants k_flip_,⊓, converted from the frequencies in the NGS results. We sought to calculate the value of the weight scores that would yield the best fit by linear regression.^25^ First, the input sequence as a character string was converted into a two-dimensional matrix (Table S7). We expressed the contribution of different bases to the overall flipping rate constant k_flip_ by equation (1), which assumed that the overall k_flip_ was the product of the weight fractions W_ij_ of the nine positions. By taking the logarithm of both sides in equation (1), the resulting equation (2) is linear with respect to all log(W_ij_ values since only one X_ij_ value is non-zero for a given value of i.

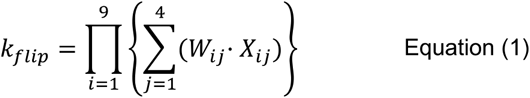

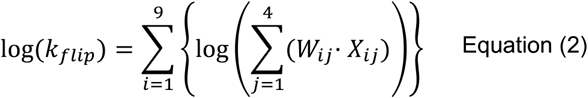

In library 2, we analyzed the 3000 most enriched sequences under selection with 1 nM Bxb1 and derived the corresponding weight scores Wij by linear regression (Fig. 3B). The W_i1_ values, corresponding to the WT reference sequence, were set to 1, which in turn produces an overall k_flip, WT_ = 1. Analyzing the fitted Wij for all position bases, we found that the weight scores of most substitutions were smaller than WT base at each position, which is consistent with the expectation that most mutations would be less efficient than WT even though the library 2 was designed to largely exclude highly conserved positions. Specifically, some substitutions resulted in moderate loss of DNA recombination function (0.1 < W < 1), including 3T, 5G, 8A, 11A, 11G, 15G, 15T, 16C, 17C, and 17G. Other substitutions such as 3A, 3C, 11C, 12T, 14A, 14G, 15A, 16G, and 16T greatly disrupted DNA recombination (W < 0.1). Notably, several substitutions – 5A, 5T, 8T, 12A, 12C, 14T, and 17A – had a W > 1, suggesting higher flipping rates than the WT sequence. To examine whether the weight scores obtained from one set of sequences can accurately predict the reaction rates for another set of sequences, we randomly split 200 sequences into a training set (140 sequences) and a validation set (60 sequences). The weight scores derived from the training sequences (Fig. 3C) were able to accurately predict the DNA recombination efficiency of the validation sequence set (Fig. 3D).

### Model evaluation

To further evaluate the model and to exclude the possibility of overfitting, we randomly selected different numbers of sequences (from 10 to 3000) to train our model, and then analyzed the variation of the weight scores and prediction errors as a function of sequence number (Fig. 4A). The prediction error for both the training and validation data sets was defined as the difference between the predicted output and the experimental output of log(k_flip_). As seen in Fig. 4B, the prediction error of the validation set was much larger than that of the training set when the total number of sequences was less than 100, suggestive of overfitting. This conclusion is also supported by the highly variable W values when the sequence number is less than 100 (Fig. 4A). As more sequences were used for model training, the prediction set error and the validation set error converged to an acceptable value (log(k_flip_) < 0.1) around 100-200 sequences (Fig. 4A,B), suggesting that this number is sufficient to accurately predict the performance of each individual sequence in a DNA library containing 250,000 variants.

**Figure 4.**
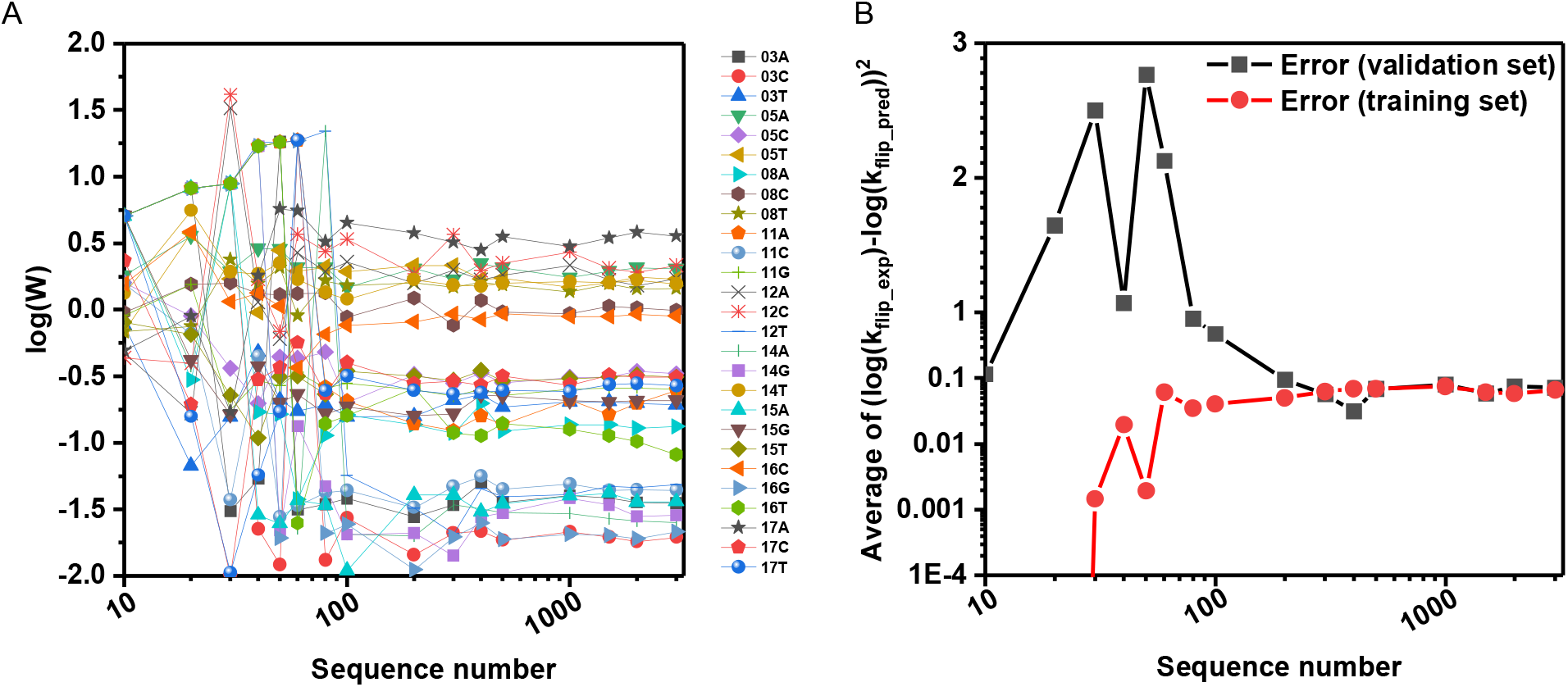
Data size sensitivity of the model. (A) Generated weight score values, W, as a function of the size of the data set (the number on the x-axis is total data set, 70% of which are training data and 30% of which are validation data). (B) Difference between predicted log(k_flip_) and experimental log(k_flip_) of the training data set (red) and validation data set (black).

When analyzing sequences selected with 1 nM Bxb1 for 5 min, we obtained weight scores that were good predictors of the overall reaction rate constant, k_flip_. Given that the specific selection conditions can influence the weight scores, as evidenced in Fig. 2D, we analyzed DNA sequences selected under different Bxb1 concentrations. In comparing the results of weight scores obtained under 1 nM, 3 nM, and 10 nM Bxb1 selection conditions, we found that for most of the positions, the values obtained under different selection conditions were similar (Fig. S4), indicating that nucleotide substitutions that are intrinsically more or less efficient than wild-type in facilitating SSR are robustly captured by our model.

The magnitude of the weight scores decreased as the concentration of Bxb1 increased from 1 nM to 3 nM to 10 nM in the selections (Fig. S4). This is not unexpected since increasing the enzyme concentration can offset biochemical advantages or disadvantages that specific nucleotide substitutions generate in enzyme-substrate affinity or catalytic turnover.

As previously noted, the logarithm of k_flip_ is proportional to the sum of the sequence weight scores in our model, which assumes independence of the contributions of individual nucleotides to the overall reaction rate constant. In protein engineering, multiple residue substitutions may have cooperative interactions or cause significant conformational changes at the binding or catalytic site; however, for the nucleotide substitutions here, significant structural changes at the protein-DNA interface are less likely to occur due to the rigid double-stranded helical structure of the DNA and the tolerance of recombinases to some sequence substitutions in their target DNA.^26^ Furthermore, based on homology modeling of the Bxb1-DNA crystal structure complex (Fig. S5), we found that the high conservation at certain nucleotide positions is due to strong recombinase interactions, both from the methyl groups on thymine and specific hydrogen bond donors and acceptors in the major groove.^21–23^ In contrast, nucleotides amenable to substitutions have long-range interactions with flexible linker loops, such as water-mediated and electrostatic interactions.^20^ At these positions, each nucleotide makes a roughly independent contribution to the long-range interaction to change the overall free energy of protein-DNA binding. Although this observation has not been reported previously for recombinase-DNA interactions, this independent base contribution has been demonstrated for other well-studied protein-DNA interactions, including transcription factor (TF)-promoter interactions, and modeled using similarly defined position weight matrices.^27,28^ Despite its predictive capability, our model alone cannot elucidate why specific nucleotide substitutions tune the reaction rate. While the model highlights specific putative interactors, a high-resolution structure of the Bxb1-DNA complex would help to mechanistically interpret the model weight scores.

### Experimental model validation and applications

To experimentally validate that the predicted k_flip_ values derived from DNA libraries can accurately predict the reaction rate of individual sequences, we tested individual attP-L variants using our *in vitro* inversion reaction assay. We cloned a panel of attP-L variants with different predicted flipping rates, including three sequences predicted to be more efficient than WT (Fig. 5A, top), and inserted them into linear DNA fragments. To test the predictive capability of the model, we added 10 nM Bxb1 recombinase to the reaction buffer and measured the percentage of DNA flipping by qPCR for each sequence after a 5-minute incubation at 30 °C. The predicted and measured flipped percentage values for these 12 sequences, including the most efficient attP-L mutant (Sequence 1), showed good correlation (Fig. 5A, bottom). We also demonstrated that the mutant sequences had a predictable range over four orders of magnitude, which provides more tunability for synthetic circuits in which SSR is a key element in the gene network.

**Figure 5:**
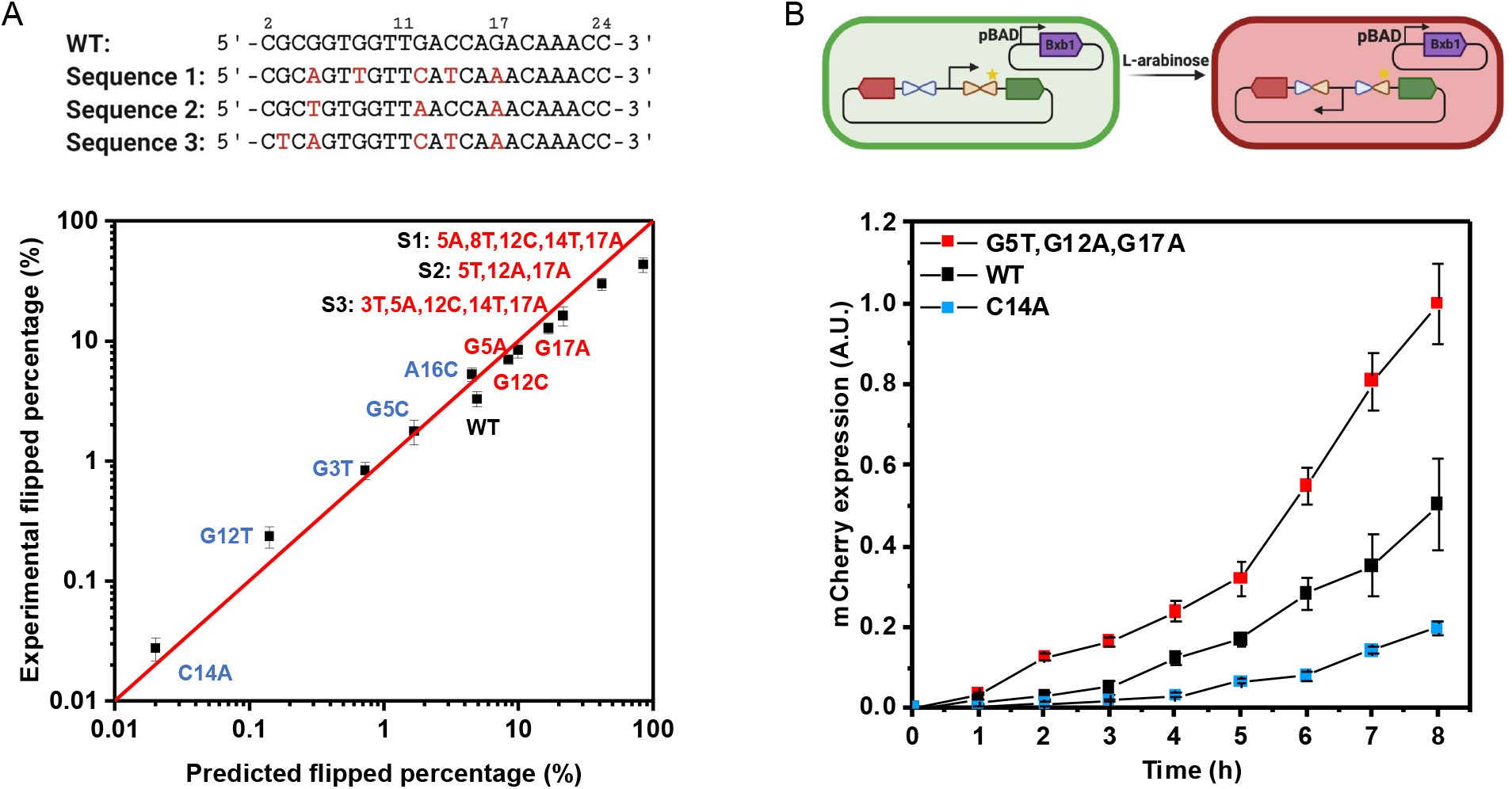
Validation of individual sequences by measuring DNA flipped percentage in reaction buffer and mCherry expression in *E. coli* cells. (A) Accurate prediction of DNA flipped percentage of multiple individual sequences with various predicted k_flip_ values. Bxb1 concentration is 10 nM and DNA fragment concentration is 1 nM. The reactions were incubated at 30 °C for 5 min and then terminated by denaturation at 80 °C for 20 min. The flipped DNA fragment concentration was measured by qPCR. Sequences 1, 2, and 3 at the top were predicted to be the most efficient variants, each incorporating multiple mutations. (B) Three sequences with significantly different DNA flipping performances (see panel A), tested in *E. coli*. Top: Donor plasmid with Bxb1 gene (purple block) and substrate plasmid are transformed into *E. coli*. DNA flipping of the promoter sequence between the GFP gene (green block) and mCherry gene (red block) gene will result in a switch from GFP expression (left) to mCherry expression (right). Bottom: The different flipping rates of these attP-L variants can be characterized by mCherry expression since it is only expressed when the promoter DNA sequence is flipped. mCherry expression level is normalized to the highest intensity of the sequence with G5T, G12A, G17A mutations at the 8-h time point. The error bars represent biological duplicates.

To further confirm that our predictions of inversion rates translate into cells, we characterized flipping dynamics of a genetic circuit element in *E. coli*. For this experiment, we transformed a Bxb1 donor plasmid and a substrate plasmid into *E. coli* to monitor the DNA flipping process (Fig. 5B, top). We inserted three different attP-L sequences into the substrate plasmid to modulate the flipping rate of the promoter by Bxb1, which was monitored by mCherry expression (Fig. 5B, bottom). The sequence with the faster predicted reaction rate than WT resulted in higher mCherry expression and the sequence with the slower predicted rate resulted in a longer lag phase and lower mCherry expression at the same growth rate. Thus, our model-guided DNA engineering approach can programmably regulate protein expression profiles in synthetic gene circuits.

These DNA attachment sequences with programmable recombination rates can serve as useful tools for kinetic control in synthetic gene circuit design. For instance, by incorporating attP variants with predictable reaction rates into a gene circuit that co-expresses two proteins, their proportions and total expression level can be rationally tuned without the need for external control. As shown in Fig. 6A, GFP and mCherry expression levels are regulated by attP1 and attP2 recombination rates, respectively. Notably, there is product inhibition in this system, as illustrated in Fig. 6B. For example, after an attP1-mediated inversion, the intact attP2 site could still bind the Bxb1 recombinase even though a recombination even is no longer possible; the same would be true for an attP1 site after an attP2-mediated inversion. These free sites serve as decoys that reduce the effective concentration of the recombinase and, depending on the k_flip_ values for the selected attP variants, this enzyme sequestration phenomenon can modulate the total protein expression. To examine these system dynamics more quantitatively, we constructed a mathematical model based on the reactions in Fig. 6B (see Supplemental Method 3) and simulated relative and total expression levels as a function of the k_flip_ values for the two attP sites (Fig. 6C). For nine selected k_flip_ combinations (numbered in Fig. 6C and shown as bar graphs in Fig. 6D, top), a range of relative and total expression levels are predicted. As expected, simulations with larger k_flip_(attP1) and k_flip_(attP2) values result in higher GFP and mCherry expression levels, respectively. More interestingly, the magnitudes of the k_flip_ values dictate the total expression level. For example, attP2 in sample 8 has a smaller recombination rate constant than that in sample 4, resulting in a smaller mCherry:GFP ratio; however, the total amount of fluorescent protein expression in sample 8 was larger than that in sample 4, due to weaker product inhibition in sample 8. These nine attP combinations were experimentally constructed and tested, showing good agreement with the model predictions in both relative and total expression levels. This example highlights the utility of differential kinetic control in gene circuit design and should have broad utility in synthetic biology to rationally tune dynamics of gene expression and memory storage.

**Figure 6:**
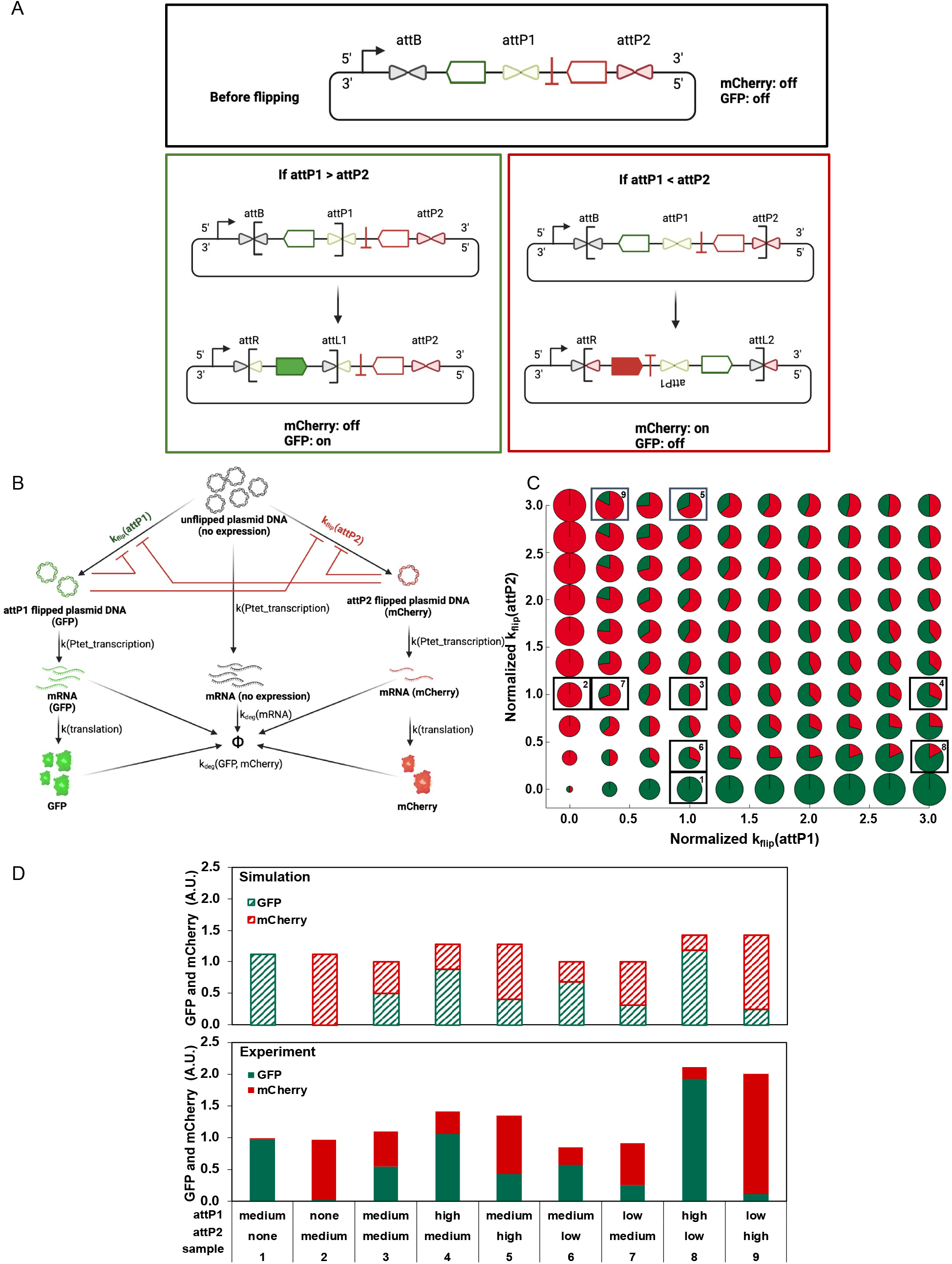
Use of attP variants with predictable DNA recombination rates to coordinate co-expression of GFP and mCherry in *E. coli*. (A) Plasmid design for GFP and mCherry co-expression. At the top is the plasmid construct before flipping. A reversed GFP gene, flanked by attB and attP1, and a reversed mCherry gene, positioned between attP1 and attP2, are placed downstream of a promoter; attP1 and attP2 are variants with different DNA recombination rates. When the recombination rate of attP1 is larger than attP2 (bottom left), the GFP gene in the bracket will be preferentially flipped and transcribed. When the recombination rate of attP2 is larger than attP1 (bottom right), the entire construct in the bracket, which includes both the GFP and mCherry genes, will be flipped; however, due to the terminator immediately after the mCherry gene, only mCherry is expressed. (B) Schematic of the GFP/mCherry co-expression network in *E. coli*, used to construct a mechanistic model (Supplemental Method 3). (C) Bubble-pie chart of simulations of the mCherry/GFP co-expression levels in *E. coli* using attP1 and attP2 variants with different DNA flipping rates. The green and red colors indicate relative proportions of GFP and mCherry expression, respectively, and the pie size indicates the total expression of fluorescent protein. The predictions tested experimentally in panel D are boxed and numbered on this pie chart. (D) Simulated (top) and experimental (bottom) GFP and mCherry expression levels for different attP1 and attP2 combinations, after 12-h incubation in M9 medium at 37 °C and normalized by cell density. Across the sample set, the low attP sequence was C14A; the medium attP sequence was WT; and the high attP sequence was G5T, G12A, G17A. The relative expression level was normalized by sample 1: WT attP1 and no attP2.

## Conclusions

Previous characterization studies of site-specific recombinases have focused on the specificity of DNA recognition,^21,22^ but quantitative analysis of the reaction rate dependence on DNA sequence has not been reported. Our qPCR-based assay is a rapid *in vitro* method for capturing initial SSR reaction rates. In addition, we developed a data-driven method to predict the relative reaction rate as a function of nucleotide sequence. Using this approach, the reaction rate can be programmed over four orders of magnitude by making rational nucleotide substitutions in the attP-L half-site sequence. By combining multiple substitutions that individually make modest improvements in the recombination efficiency, we created a new attP-L sequence that confers a 10-fold increase in initial reaction rate over WT (Fig. 5A). This sequence (S1: 5’-CGCAGTTGTTCATCAAACAAACC-3’) should be useful in improving Bxb1 recombination efficiency across a range of synthetic biology and cell engineering applications. Moreover, the use of model-guided design to rationally tune Bxb1 recombination reaction rates enables differential kinetic control of multiple processes by the same enzyme, which can allow for stoichiometric control of the expression of multiple proteins, as shown here, but also temporal ordering of events in a synthetic genetic program without the need for multiple enzymes.

This quantitative data-driven approach can be extended to other recombinases and DNA or RNA modifying enzymes for which an appropriate selection system can be developed. Importantly, this method can be applied without a detailed mechanistic or structural understanding of the complex interaction between protein and DNA. In fact, the computed weight scores, which are derived from kinetic experiments, can reveal indirect long-range interactions that are important for the observed dynamics but may not be captured by static crystal structures. Deriving the corresponding weight score matrices for other such enzymes can therefore enhance our mechanistic understanding of their function, further broaden the genetic editing toolbox, and enhance the ability to design tunable artificial circuits.

## Methods

### General methods

All cloning was performed in *E. coli* strain XL1-blue (Agilent, 200130); BL21(DE3) competent cells were used for recombinant Bxb1 protein expression; and *E. coli* strain TOP10 pro (F’[lacIq Tn10(tetR)] mcrA Δ(mrr-hsdRMS-mcrBC) φ80lacZΔM15 ΔlacX74 deoR nupG recA1 araD139 Δ(ara-leu)7697 galU galK rpsL(StrR) endA1 λ-, a generous gift from Dr. James J. Collins) was used for all experimental characterization. *E. coli* cells were grown in LB medium (Fisher Scientific, DF0446173), SOB medium (Teknova, S0210), or M9 medium containing M9 salt (Sigma-Aldrich, M6030), 0.4 % glycerol (Sigma-Aldrich, G7757), 0.2 % casamino acids (MP Biomedicals, 113060012), 2 mM MgSO4 (Fisher Scientific, BP213), 0.1 mM CaCl2 (Fisher Scientific, BP510), and 0.34 g/L thiamine hydrochloride (Sigma-Aldrich, T4625). Antibiotic concentrations of 100 μg/ml carbenicillin (Teknova, C2105) and 15 μg/ml chloramphenicol (Sigma-Aldrich, C0378) were used to maintain plasmids in *E. coli*. Isopropyl β-D-1-thiogalactopyranoside (IPTG, Millipore Sigma, 420322), anhydrotetracycline hydrochloride (aTc, AdipoGen, CDX-A0197-M500), and L-arabinose (Sigma-Aldrich, A3256) were used to induce gene expression.

Oligonucleotides were purchased from Integrated DNA Technology (IDT). Phusion High-Fidelity Polymerase from New England Biolabs (NEB, M0530) was used for all PCR amplifications. PowerUp SYBR Green Master Mix (Thermo Fisher Scientific, A25741) was used for qPCR. T4 ligase and restriction enzymes for gene cloning were purchased from NEB. Plasmid and DNA fragments were purified using kits from Qiagen according to manufacturer’s instructions. The sequences were verified by Sanger sequencing by ACGT, INC.

### Construction of the attP-L library

DNA oligonucleotides containing the randomized attP-L DNA sequences were used as the reverse primer for PCR amplification. After purification, the DNA fragment with the attP-L variants at the end was digested by HindIII-HF and ligated to another DNA fragment with the complementary HindIII-HF sticky end. Thus, the attP-L library was in the middle of a linear DNA fragment after ligation. The ligation product was purified by gel extraction and was used as the substrate for library selection. Sequence information and the DNA oligos can be found in Table S1. The quality of the naïve constructed library was verified by NGS, as shown in Table S5 and S6.

### Expression and purification of Bxb1 recombinase

The Bxb1 gene was cloned from a plasmid (Addgene #123132) and inserted into the expression vector pET22, which appends a C-terminal (His)_6_ tag. Primers are listed in Table S1. The plasmid pET22-Bxb1was transformed into BL21(DE3) cells. A sequence-verified colony was grown in 5 ml LB medium overnight at 37 °C. Then the overnight cell culture was diluted to OD_600_ = 0.05 in 2x YT medium and grown at 30 °C until OD_600_ = 0.6. To induce Bxb1 expression, 0.4 mM IPTG was added and the incubation temperature was decreased to 16 °C. After a 15-h incubation, the cells were harvest by centrifuging the cell culture at 4000*g* for 30 min.

The cell pellet was suspended in a lysis buffer and lysed by sonication. The cell lysate was then centrifuged to remove cell debris. The purification was performed using Ni-NTA resin from Qiagen. After equilibrating the 1 ml 50% resin using 5 ml lysis buffer (50 mM Tris-HCl, pH 8, 300 mM NaCl, 10 mM imidazole, 1 mM DTT), the cell lysate was loaded onto the column. The resin was then washed with 25 ml wash buffer 1 (50 mM Tris-HCl, pH 8, 300 mM NaCl, 10 mM imidazole, 1 mM DTT) and 2.5 ml wash buffer 2 (50 mM Tris-HCl, pH 8, 800 mM NaCl, 65 mM imidazole, 1 mM DTT). Then the protein was diluted by a 2 ml elution buffer (50 mM Tris-HCl, pH 8, 800 mM NaCl, 200 mM imidazole, 1mM DTT). Finally, the protein was rebuffered and concentrated via centrifugal ultrafiltration using a column with 10-kDa cutoff (Millipore). The purified protein was stored in storage buffer (50 mM Tris-HCl, pH 8, 300 mM NaCl, 1 mM DTT, 1 mM EDTA, and 50% glycerol) at −20 °C. The protein size was verified by SDS-PAGE and the concentration was quantified by BCA assay.

### In vitro recombination assays

The DNA recombination reactions were performed in PCR tubes in a reaction volume of 50 μl. Purified linear DNA fragments containing the Bxb1 attachment DNA sites in the middle were used as substrate. The reaction buffer consisted of 50 mM NaCl, 10 mM Tris-HCl, pH 7.9, 10 mM MgCl2, and 100 μg/ml BSA. To test various Bxb1 concentrations, the 1 μg/μl concentrated Bxb1 stock was serially diluted to 10 times the desired final concentration using the same storage buffer, so that 5 μl 10x Bxb1 enzyme could be added to a 50 μl reaction. The reaction was incubated at 30 °C and terminated by heating at 80 °C for 20 min.

After recombination, the percentage of flipped DNA substrate was quantified using qPCR to measure the number of flipped molecules. The recombination mixture was diluted using Milli-Q water so that the total DNA concentration was 10^9^ copies/μL. In a 20 μL qPCR reaction, 0.5 μM primer, 1 μL of diluted recombinant reaction mixture (10^9^ copies of total DNA), and 10 μl of PowerUp SYBR Green master mix were added. PCR cycles were set up as follows: preincubation at 95 °C for 2 min, then 40 cycles were repeated with denaturation at 95 °C for 15 s, annealing at 60 °C for 15 s, and extension at 72 °C for 15 s. Dissociation curve analysis was then performed to test qPCR sensitivity. The same peak position in the dissociation curves indicated that the unflipped DNA substrate does not interfere with the PCR reaction. Through appropriate primer design, qPCR selectively amplified only the flipped DNA template. The primer sequences are listed in Table S1.

### Amplicon preparation for sequencing

Two rounds of PCR were performed to barcode the selected DNA libraries. In the first round, a forward primer and reverse primer were used to introduce a constant region that would serve as an annealing site in the second round of PCR. In the second round, the Nextera DNA CD index was used as the primers. The 5’ end of the index was used for binding between the amplicon and the sequencing flow cell, the 3’ end was used for annealing to the PCR product from the first round, and the middle 8 bases, i5 or i7, were used for barcoding different samples. DNA samples were quantified with the Quant-iT PicoGreen dsDNA detection kit (Thermo Fisher Scientific) according to the manufacturer’s instructions. Sequencing was performed on an Illumina MiniSeq using a paired-end read kit following the manufacturer’s instructions.

### NGS data analysis

The FASTQ sequencing results generated from the MiniSeq were written to .csv files in Matlab (Mathworks). To refine the results, reads that did not match at consistent positions due to sequencing errors were filtered out. For the remaining sequences, duplicate sequences were removed, and only unique sequences were listed. For each distinct sequence, its frequency of occurrence in the total number of refined sequences was counted. In addition, the enrichment of each of the four nucleotides at each randomized position was also calculated.

### Validation in E. coli

The high-copy-number pUC vector was used for the construction of the GFP/mCherry reporter plasmid and the low copy number p15A_PBAD vector was used for the construction of the Bxb1 donor plasmid, using restriction enzyme digestion and ligation. The primers, plasmids, and strains used are listed in Tables S1, S2, and S3, respectively.

Each plasmid (100 ng) was transformed into 50 μL TOP10 pro competent cells at 42 °C for 45 s by heat shock. Then, 950 μL of pre-warmed SOB medium was added, followed by incubation at 37 °C and 200 rpm for 1 h. After recovery, the cell cultures were centrifuged at 3000*g* for 90 s and resuspended in M9 minimal medium with 50 μg/ml carbenicillin, 15 μg/ml chloramphenicol, and 0.1% L-arabinose inducer for Bxb1 expression. Cells were aliquoted into 96-well plates (200 μL/well) and incubated in a plate reader (Cytation 3 image reader) at 30°C and a shaking speed of 807 rpm. Cell growth (OD_600_), GFP expression (ex/em: 488 nm/509 nm), and mCherry expression (ex/em: 584 nm/610 nm) were monitored inline for 8 h on the plate reader. For each transformation, two biological replicates and three technical replicates were performed. Strains transformed with different reporter plasmids had similar growth rates (Fig. S4). GFP and mCherry expression levels were normalized to OD_600_ and subtracted from the basal fluorescence readings of the negative control strain (TOP10 pro cells without plasmid transformation).

## Supporting Information

Table S1: Oligonucleotides used in this study

Table S2: Plasmids used in this study

Table S3: Strains used in this study

Table S4. Sequences of library 1 and library 2

Table S5. Nucleotide enrichment of library 1 after selection with different Bxb1 concentrations Table S6. Nucleotide enrichment of library 2 after selection with different Bxb1 concentrations Table S7. Conversion of DNA sequences into binary matrices

Figure S1: *In vivo* selection in *E. coli*

Figure S2: Wild-type DNA inversion time course quantified by qPCR and gel electrophoresis Figure S3: Correlation between normalized frequency and normalized DNA flipping rate constant k_flip_

Figure S4: Comparison of weight scores for different Bxb1 concentrations

Figure S5: Posited interactions between Bxb1 and attP-L based on generated weight scores and homology modeling

Supplemental Method 1: *In vivo* selection using *E. coli*

Supplemental Method 2: High-throughput reaction rate characterization *in vitro* using NGS Supplemental Method 3: Mathematical simulation of GFP and mCherry co-expression in *E. coli*

## Notes

The authors declare no competing financial interest.

## Acknowledgements

We thank Victor Garcia for assistance with next-generation sequencing, and Ayako Ohoka and Jennifer Kang for helpful discussions. This work was supported by the National Institutes of Health (R01DK114453 to S.M.A. and C.A.S.).

## Notes

### Competing Interest Statement

The authors have declared no competing interest.

