## Supplementary Information for "Model-guided engineering of DNA sequences with predictable site-specific recombination rates"

### Supplemental Tables

**Table S1.** Oligonucleotides used in this study

| Purpose | Name | Sequence (5' → 3') |
|---|---|---|
| Linear DNA library | F-library upstream | AGGCCTTGGAGCCGTACATG |
| | F-library downstream | CCCGGGAAGCTTTTTAAGAAGGAGATATACATATGAGCAAAGGAGAAGAACTTTT |
| | R-library downstream | TCCGTTTGTAGCATCACC |
| | R-library1 upstream | CTTAAAAAGCTTTCGT**GGTTTGNNNNNNNNNNCACCGCGG**TCTCAGTGGT |
| | R-library2 upstream | CTTAAAAAGCTTTCGT**GGTTTNTNNNNTNNACNACNGNGG**TCTCAGTGGTGTACGG |
| | R-WT upstream | CTTAAAAAGCTTTCGT**GGTTTGTCTGGTCAACCACCGCGG**TCTCAGTGGT |
| Bxb1 purification plasmid: pET22-bxb1 | F-bxb1 | GATATACATATGGTGAGAGCCCTGGTAG |
| | R-bxb1 | GTGGTGGGTCTCCTCGAGCGACATCCCGGTGTGTAG |
| MiniSeq amplicon preparation | F-first step | TCGTCGGCAGCGTCAGATGTGTATAAGAGACAGTGATCCTGACGACGGAGA |
| | R-first step | GTCTCGTGGGCTCGGAGATGTGTATAAGAGACAGAAAAAGCTTTCGTGGTTT |
| qPCR quantification | F-qPCR | TTGTGCCCATTAACATCACC |
| | R-qPCR | ATGATCCTGACGACGGAGA |
| Variants for *in vitro* validation | R-C14A | CTTAAAAAGCTTTCGT**GGTTTGTCTGTTCAACCACCGCGG**TCTCAGTGGT |
| | R-G5C | CTTAAAAAGCTTTCGT**GGTTTGTCTGGTCAACCACGGCGG**TCTCAGTGGT |
| | R-A16C | CTTAAAAAGCTTTCGT**GGTTTGTCGGGTCAACCACCGCGG**TCTCAGTGGT |
| | R-G12C | CTTAAAAAGCTTTCGT**GGTTTGTCTGGTGAACCACCGCGG**TCTCAGTGGT |
| | R-G5A | CTTAAAAAGCTTTCGT**GGTTTGTCTGGTCAACCACTGCGG**TCTCAGTGGT |
| | R-G17A | CTTAAAAAGCTTTCGT**GGTTTGTTTGGTCAACCACCGCGG**TCTCAGTGGT |
| | R-3T, 5A, 12C, 14T, 17A | CTTAAAAAGCTTTCGT**GGTTTGTTTGATGAACCACTGTGG**TCTCAGTGGT |
| | R-5T, 12A, 17A | CTTAAAAAGCTTTCGT**GGTTTGTTTGGTTAACCACAGCGG**TCTCAGTGGT |
| | R-5A, 8T, 12C, 14T, 17A | CTTAAAAAGCTTTCGT**GGTTTGTTTGATGAACAACTGCGG**TCTCAGTGGT |
| pUC-gfp/mcherry for *in vivo* validation | F-mCherry | CCCGGGTCTAGAGCGAGTCAG |
| | R-mCherry | GGGCCCGGATCCTTTAAGAAGGAGAT |
| | F-attB | GGGCCCGGATCCTCGGCCGGCTTGTC |

| | | |
|---|---|---|
| | R-attP | CCCGGGAAGCTTTCGTGGTTTGTCTGGTCAACCACCGCGGTCTCAGTGGTGTACGGTACAAACCCGCTAGCAGCTGTTTCCTGTGTGAAATTGTTATCCG |
| | F-GFP | CCCGGGAAGCTTTTTAAGAAGGAGATATACATATGAGCAAAGGAGAAGAACTTTT |
| | R-GFP | GGGCCCGGTACCTAAAACGACGGCCAGTGAATTCGAGCTCCAAAAAAACCCCTCAAGACCC |
| | F-pUC vector | GGGCCCGGTACCCAACGTCGTGACTGGG |
| | R-pUC vector | CCCGGGTCTAGATGCGCTCGGTCGTTC |
| Bxb1 donor plasmid for *in vivo* validation: p15A-$P_{BAD}$-bxb1 | F-PBAD | ACCTGGTGCACGGCAGATACACTTGCTG |
| | R-PBAD | CTTGAGGAATTCTTGAAGACGAAAGGGC |
| | F-bxb1 (PBAD) | TCGTCTTCAAGAATTCCTCAAGAAGATCCTTTGATC |
| | R-bxb1 (PBAD) | TCTGCCGTGCACCAGGTGTCTAGAAGAGACAC |
| KanR/CmR reporter plasmid for preliminary selection *in vivo*: p15A-$P_{cons}$-kanR/cmR | F-KanR | GGTGTTGGTACCTCGGCCGGCTTGTCGACGACGGCGGTCTCCGTCGTCAGGATCATCCGGGCAGGAGATATACATATGATGAGCCATATTCAACGG |
| | R-KanR | GCGTAAAAGCTTTTAGAAAAACTCATCGAGC |
| | F-CmR | TTCTAAAAGCTTTTACGCCCCGCCCTGCCAC |
| | R-CmR | CAATTGGGATCCAGGAGATATACATATGATGGAGAAAAAAATCACTGG |
| Bxb1 donor plasmid for preliminary selection *in vivo*: pUC-bxb1 | F-bxb1 (pUC) | TCCCCGGGTACCGTGAGAGCCCTGGTAG |
| | R-bxb1 (pUC) | TGCACCATATGCTTACGACATCCCGGTGTGTAG |

**Table S2.** Plasmids used in this study

| Plasmid | Name | Origin of replication | Promoter | Antibiotic resistance |
|---|---|---|---|---|
| Bxb1 donor for *in vivo* validation | p15A-$P_{BAD}$-bxb1 | p15A (low copy) | Arabinose promoter | Chloramphenicol |
| Bxb1 donor for *in vivo* preliminary selection | pUC19-$P_{BAD}$-bxb1 | pUC19 (high copy) | Arabinose promoter | Carbenicillin |
| Bxb1 purification | pET22-$P_{T7}$-bxb1 | pET22 (medium copy) | T7 promoter | Carbenicillin |
| GFP/mCherry reporter for *in vivo* validation | pUC19-attB-$P_{Lac}$-attP-gfp/mcherry | pUC19 (high copy) | Lactose promoter | Carbenicillin |
| KanR/CmR reporter for *in vivo* preliminary selection | p15A-attB-$P_{cons}$-attP-kanR/cmR | p15A (low copy) | Constitutive promoter | Kanamycin or Chloramphenicol |

**Table S3.** Strains used in this study

| **Purpose** | **Host cell** | **Plasmid 1** | **Plasmid 2** |
|---|---|---|---|
| *In vivo* validation of attP variants | TOP10 pro | p15A-$P_{BAD}$-bxb1 | pUC19-attB-$P_{Lac}$-attP-gfp/mcherry |
| Negative control for *in vivo* validation of attP variants | TOP10 pro | -- | pUC19-attB-$P_{Lac}$-attP-gfp/mcherry |
| Preliminary library 1 selection *in vivo* | TOP10 pro | pUC19-$P_{BAD}$-bxb1 | p15A-attB-$P_{cons}$-attP-kanR/cmR |
| Bxb1 protein purification | BL21(DE3) | pET22-$P_{T7}$-bxb1 | -- |

**Table S4.** Sequences of library 1 and library 2

| | **Position** | | | | | | | | | | | | | | | | | | | | | | | |
|---|---|---|---|---|---|---|---|---|---|---|---|---|---|---|---|---|---|---|---|---|---|---|---|---|
| | 1 | 2 | 3 | 4 | 5 | 6 | 7 | 8 | 9 | 10 | 11 | 12 | 13 | 14 | 15 | 16 | 17 | 18 | 19 | 20 | 21 | 22 | 23 | 24 |
| WT | C | C | G | C | G | G | T | G | G | T | T | G | A | C | C | A | G | A | C | A | A | A | C | C |
| Library 1 | C | C | G | C | G | G | T | G | N | N | N | N | N | N | N | N | N | N | C | A | A | A | C | C |
| Library 2 | C | C | N | C | N | G | T | N | G | T | N | N | A | N | N | N | N | A | N | A | A | A | C | C |

**Table S5.** Nucleotide enrichment of library 1 after selection with different Bxb1 concentrations

| Selection condition | Position | 9 | 10 | 11 | 12 | 13 | 14 | 15 | 16 | 17 | 18 |
|---|---|---|---|---|---|---|---|---|---|---|---|
| | WT | G | T | T | G | A | C | C | A | G | A |
| Naïve library | A | 0.25 | 0.25 | 0.23 | 0.25 | 0.25 | 0.23 | 0.24 | 0.22 | 0.23 | 0.28 |
| | C | 0.22 | 0.27 | 0.28 | 0.26 | 0.29 | 0.27 | 0.26 | 0.27 | 0.28 | 0.30 |
| | G | 0.28 | 0.24 | 0.23 | 0.25 | 0.21 | 0.25 | 0.26 | 0.26 | 0.26 | 0.24 |
| | T | 0.25 | 0.25 | 0.26 | 0.24 | 0.25 | 0.25 | 0.24 | 0.23 | 0.23 | 0.18 |
| 300 nM, 1h | A | 0.27 | 0.14 | 0.18 | 0.24 | **0.28** | 0.15 | 0.16 | **0.25** | 0.27 | **0.35** |
| | C | 0.17 | 0.35 | 0.37 | 0.39 | 0.31 | **0.47** | **0.46** | 0.37 | 0.35 | 0.31 |
| | G | **0.42** | 0.07 | 0.21 | **0.21** | 0.22 | 0.15 | 0.22 | 0.22 | **0.20** | 0.24 |
| | T | 0.14 | **0.45** | **0.24** | 0.16 | 0.20 | 0.22 | 0.17 | 0.17 | 0.18 | 0.10 |
| 300 nM, 5min | A | 0.24 | 0.09 | 0.17 | 0.23 | **0.29** | 0.11 | 0.13 | **0.26** | 0.28 | **0.38** |
| | C | 0.12 | 0.26 | 0.34 | 0.40 | 0.29 | **0.53** | **0.49** | 0.36 | 0.33 | 0.29 |
| | G | **0.52** | 0.05 | 0.23 | **0.21** | 0.22 | 0.13 | 0.24 | 0.23 | **0.21** | 0.24 |
| | T | 0.12 | **0.60** | **0.26** | 0.15 | 0.20 | 0.23 | 0.14 | 0.15 | 0.19 | 0.09 |
| 100 nM, 5min | A | 0.23 | 0.08 | 0.17 | 0.24 | **0.29** | 0.11 | 0.12 | **0.27** | 0.30 | **0.44** |
| | C | 0.11 | 0.24 | 0.34 | 0.39 | 0.29 | **0.51** | **0.49** | 0.36 | 0.31 | 0.25 |
| | G | **0.55** | 0.04 | 0.23 | **0.22** | 0.21 | 0.13 | 0.24 | 0.22 | **0.22** | 0.24 |
| | T | 0.11 | **0.64** | **0.26** | 0.15 | 0.21 | 0.24 | 0.15 | 0.15 | 0.17 | 0.07 |
| 50 nM, 5min | A | 0.22 | 0.07 | 0.16 | 0.25 | **0.30** | 0.11 | 0.12 | **0.28** | 0.30 | **0.47** |
| | C | 0.09 | 0.22 | 0.33 | 0.38 | 0.28 | **0.51** | **0.49** | 0.35 | 0.31 | 0.24 |
| | G | **0.59** | 0.04 | 0.24 | **0.23** | 0.21 | 0.14 | 0.24 | 0.22 | **0.22** | 0.23 |
| | T | 0.10 | **0.67** | **0.27** | 0.14 | 0.20 | 0.24 | 0.15 | 0.15 | 0.16 | 0.06 |
| 25 nM, 5min | A | 0.18 | 0.06 | 0.15 | 0.26 | **0.31** | 0.12 | 0.13 | **0.32** | 0.32 | **0.52** |
| | C | 0.07 | 0.17 | 0.32 | 0.36 | 0.27 | **0.50** | **0.49** | 0.34 | 0.28 | 0.21 |
| | G | **0.67** | 0.03 | 0.24 | **0.24** | 0.22 | 0.14 | 0.23 | 0.20 | **0.24** | 0.21 |
| | T | 0.09 | **0.74** | **0.29** | 0.14 | 0.19 | 0.25 | 0.16 | 0.14 | 0.15 | 0.05 |
| 10 nM, 5min | A | 0.19 | 0.06 | 0.15 | 0.26 | **0.32** | 0.13 | 0.13 | **0.33** | 0.32 | **0.52** |
| | C | 0.08 | 0.18 | 0.32 | 0.35 | 0.26 | **0.49** | **0.49** | 0.33 | 0.28 | 0.22 |
| | G | **0.65** | 0.03 | 0.24 | **0.25** | 0.21 | 0.14 | 0.22 | 0.20 | **0.23** | 0.21 |
| | T | 0.09 | **0.72** | **0.30** | 0.14 | 0.20 | 0.24 | 0.16 | 0.14 | 0.17 | 0.06 |
| 5 nM, 5min | A | 0.18 | 0.08 | 0.15 | 0.23 | **0.34** | 0.12 | 0.13 | **0.32** | 0.28 | **0.52** |
| | C | 0.10 | 0.20 | 0.30 | 0.35 | 0.26 | **0.49** | **0.52** | 0.32 | 0.22 | 0.22 |
| | G | **0.62** | 0.04 | 0.20 | **0.25** | 0.21 | 0.13 | 0.20 | 0.21 | **0.24** | 0.20 |
| | T | 0.10 | **0.67** | **0.35** | 0.14 | 0.19 | 0.26 | 0.16 | 0.15 | 0.16 | 0.06 |
| 4 nM, 5min | A | 0.17 | 0.06 | 0.14 | 0.24 | **0.37** | 0.12 | 0.13 | **0.34** | 0.33 | **0.56** |
| | C | 0.07 | 0.16 | 0.29 | 0.35 | 0.26 | **0.48** | **0.52** | 0.33 | 0.26 | 0.19 |
| | G | **0.69** | 0.03 | 0.22 | **0.25** | 0.19 | 0.13 | 0.19 | 0.21 | **0.22** | 0.19 |
| | T | 0.07 | **0.75** | **0.35** | 0.16 | 0.18 | 0.27 | 0.16 | 0.12 | 0.19 | 0.05 |
| 3 nM, 5min | A | 0.19 | 0.07 | 0.14 | 0.27 | **0.39** | 0.15 | 0.15 | **0.34** | 0.35 | **0.49** |
| | C | 0.10 | 0.19 | 0.29 | 0.37 | 0.24 | **0.50** | **0.47** | 0.33 | 0.29 | 0.22 |
| | G | **0.63** | 0.04 | 0.21 | **0.24** | 0.20 | 0.14 | 0.21 | 0.19 | **0.21** | 0.22 |
| | T | 0.08 | **0.70** | **0.36** | 0.14 | 0.18 | 0.31 | 0.17 | 0.14 | 0.15 | 0.07 |

**Table S6.** Nucleotide enrichment of library 2 after selection with different Bxb1 concentrations

| Selection condition | Position | 3 | 5 | 8 | 11 | 12 | 14 | 15 | 16 | 17 | 19 |
|---|---|---|---|---|---|---|---|---|---|---|---|
| | WT | G | G | G | T | G | C | C | A | G | C |
| Naïve library | A | 0.23 | 0.25 | 0.23 | 0.25 | 0.24 | 0.23 | 0.24 | 0.22 | 0.23 | 0.23 |
| | C | 0.28 | 0.27 | 0.28 | 0.26 | 0.24 | 0.27 | 0.26 | 0.27 | 0.28 | 0.30 |
| | G | 0.24 | 0.24 | 0.23 | 0.25 | 0.21 | 0.25 | 0.26 | 0.26 | 0.26 | 0.24 |
| | T | 0.25 | 0.25 | 0.26 | 0.24 | 0.24 | 0.25 | 0.24 | 0.23 | 0.23 | 0.22 |
| 300 nM, 1h | A | 0.26 | 0.24 | 0.23 | 0.23 | 0.23 | 0.11 | 0.14 | **0.22** | 0.23 | 0.06 |
| | C | 0.26 | 0.23 | 0.28 | 0.29 | 0.39 | **0.47** | **0.40** | 0.30 | 0.29 | **0.80** |
| | G | **0.26** | **0.25** | **0.17** | 0.22 | **0.21** | 0.16 | 0.28 | 0.27 | **0.28** | 0.08 |
| | T | 0.23 | 0.29 | 0.33 | **0.25** | 0.17 | 0.26 | 0.19 | 0.21 | 0.21 | 0.06 |
| 300 nM, 5min | A | 0.22 | 0.26 | 0.21 | 0.22 | 0.23 | 0.06 | 0.10 | **0.23** | 0.25 | 0.02 |
| | C | 0.21 | 0.22 | 0.28 | 0.28 | 0.41 | **0.54** | **0.42** | 0.29 | 0.26 | **0.91** |
| | G | **0.30** | **0.24** | **0.16** | 0.24 | **0.21** | 0.11 | 0.33 | 0.29 | **0.28** | 0.04 |
| | T | 0.28 | 0.28 | 0.35 | **0.27** | 0.15 | 0.28 | 0.16 | 0.20 | 0.22 | 0.02 |
| 100 nM, 5min | A | 0.21 | 0.26 | 0.20 | 0.21 | 0.22 | 0.06 | 0.10 | **0.23** | 0.25 | 0.02 |
| | C | 0.20 | 0.22 | 0.27 | 0.28 | 0.41 | **0.54** | **0.42** | 0.29 | 0.26 | **0.94** |
| | G | **0.31** | **0.24** | **0.16** | 0.24 | **0.21** | 0.11 | 0.33 | 0.29 | **0.28** | 0.03 |
| | T | 0.28 | 0.28 | 0.37 | **0.28** | 0.15 | 0.29 | 0.16 | 0.19 | 0.21 | 0.02 |
| 50 nM, 5min | A | 0.19 | 0.26 | 0.19 | 0.20 | 0.23 | 0.05 | 0.09 | **0.22** | 0.25 | 0.01 |
| | C | 0.19 | 0.22 | 0.28 | 0.28 | 0.40 | **0.55** | **0.42** | 0.29 | 0.28 | **0.96** |
| | G | **0.33** | **0.24** | **0.16** | 0.24 | **0.22** | 0.11 | 0.33 | 0.30 | **0.27** | 0.02 |
| | T | 0.28 | 0.28 | 0.37 | **0.28** | 0.15 | 0.28 | 0.16 | 0.19 | 0.20 | 0.01 |
| 25 nM, 5min | A | 0.16 | 0.27 | 0.18 | 0.18 | 0.22 | 0.06 | 0.10 | **0.25** | 0.28 | 0.01 |
| | C | 0.17 | 0.20 | 0.25 | 0.28 | 0.40 | **0.53** | **0.42** | 0.29 | 0.25 | **0.97** |
| | G | **0.38** | **0.21** | **0.15** | 0.25 | **0.22** | 0.12 | 0.33 | 0.29 | **0.28** | 0.01 |
| | T | 0.30 | 0.31 | 0.42 | **0.29** | 0.15 | 0.29 | 0.16 | 0.18 | 0.20 | 0.01 |
| 10 nM, 5min | A | 0.16 | 0.28 | 0.17 | 0.18 | 0.22 | 0.06 | 0.10 | **0.25** | 0.28 | 0.01 |
| | C | 0.17 | 0.19 | 0.24 | 0.27 | 0.41 | **0.53** | **0.42** | 0.29 | 0.25 | **0.96** |
| | G | **0.38** | **0.21** | **0.15** | 0.25 | **0.23** | 0.12 | 0.33 | 0.28 | **0.28** | 0.01 |
| | T | 0.30 | 0.32 | 0.44 | **0.30** | 0.15 | 0.29 | 0.15 | 0.17 | 0.19 | 0.01 |
| 5 nM, 5min | A | 0.17 | 0.26 | 0.18 | 0.18 | 0.22 | 0.06 | 0.09 | **0.24** | 0.27 | 0.01 |
| | C | 0.17 | 0.22 | 0.25 | 0.27 | 0.40 | **0.53** | **0.42** | 0.29 | 0.27 | **0.95** |
| | G | **0.36** | **0.21** | **0.15** | 0.24 | **0.23** | 0.12 | 0.32 | 0.29 | **0.27** | 0.02 |
| | T | 0.30 | 0.31 | 0.42 | **0.30** | 0.15 | 0.28 | 0.16 | 0.18 | 0.19 | 0.02 |
| 4 nM, 5min | A | 0.13 | 0.27 | 0.15 | 0.15 | 0.20 | 0.07 | 0.10 | **0.31** | 0.35 | 0.01 |
| | C | 0.16 | 0.18 | 0.19 | 0.26 | 0.41 | **0.51** | **0.43** | 0.27 | 0.22 | **0.96** |
| | G | **0.42** | **0.17** | **0.13** | 0.25 | **0.25** | 0.13 | 0.32 | 0.27 | **0.26** | 0.01 |
| | T | 0.28 | 0.38 | 0.53 | **0.34** | 0.14 | 0.28 | 0.16 | 0.15 | 0.17 | 0.02 |
| 3 nM, 5min | A | 0.13 | 0.27 | 0.15 | 0.14 | 0.20 | 0.07 | 0.10 | **0.31** | 0.35 | 0.01 |
| | C | 0.17 | 0.17 | 0.19 | 0.26 | 0.41 | **0.52** | **0.42** | 0.27 | 0.22 | **0.96** |
| | G | **0.43** | **0.17** | **0.13** | 0.25 | **0.25** | 0.13 | 0.32 | 0.27 | **0.26** | 0.01 |
| | T | 0.28 | 0.39 | 0.53 | **0.35** | 0.14 | 0.28 | 0.16 | 0.15 | 0.17 | 0.01 |
| 2 nM, 5min | A | 0.08 | 0.32 | 0.11 | 0.14 | 0.20 | 0.06 | 0.11 | **0.29** | 0.32 | 0.01 |
| | C | 0.09 | 0.14 | 0.20 | 0.25 | 0.39 | **0.54** | **0.42** | 0.31 | 0.24 | **0.97** |
| | G | **0.57** | **0.12** | **0.13** | 0.29 | **0.27** | 0.12 | 0.33 | 0.27 | **0.27** | 0.01 |
| | T | 0.25 | 0.42 | 0.56 | **0.32** | 0.14 | 0.27 | 0.14 | 0.13 | 0.16 | 0.01 |
| 1 nM, 5min | A | 0.10 | 0.24 | 0.09 | 0.15 | 0.18 | 0.09 | 0.14 | **0.32** | 0.29 | 0.01 |
| | C | 0.08 | 0.28 | 0.22 | 0.24 | 0.40 | **0.53** | **0.41** | 0.29 | 0.30 | **0.97** |
| | G | **0.57** | **0.13** | **0.15** | 0.28 | **0.27** | 0.14 | 0.30 | 0.21 | **0.23** | 0.01 |
| | T | 0.25 | 0.35 | 0.54 | **0.33** | 0.14 | 0.24 | 0.14 | 0.17 | 0.19 | 0.01 |

**Table S7.** Conversion of DNA sequences into binary matrices

| Position | 3 | 5 | 8 | 11 | 12 | 14 | 15 | 16 | 17 |
|---|---|---|---|---|---|---|---|---|---|
| **1 (WT reference)** | G | G | G | T | G | C | C | A | G |
| **2** | A | A | A | A | A | A | A | C | A |
| **3** | C | C | C | C | C | G | G | G | C |
| **4** | T | T | T | G | T | T | T | T | T |

*Example matrix for sequence GAGCGACGT:*

| Sequence | G | A | G | C | G | A | C | G | T |
|---|---|---|---|---|---|---|---|---|---|
| $X_{ij}$ | i=1 | i=2 | i=3 | i=4 | i=5 | i=6 | i=7 | i=8 | i=9 |
| **j=1** | 1 | 0 | 1 | 0 | 1 | 0 | 1 | 0 | 0 |
| **j=2** | 0 | 1 | 0 | 0 | 0 | 1 | 0 | 0 | 0 |
| **j=3** | 0 | 0 | 0 | 1 | 0 | 0 | 0 | 1 | 0 |
| **j=4** | 0 | 0 | 0 | 0 | 0 | 0 | 0 | 0 | 1 |

### Supplemental Figures

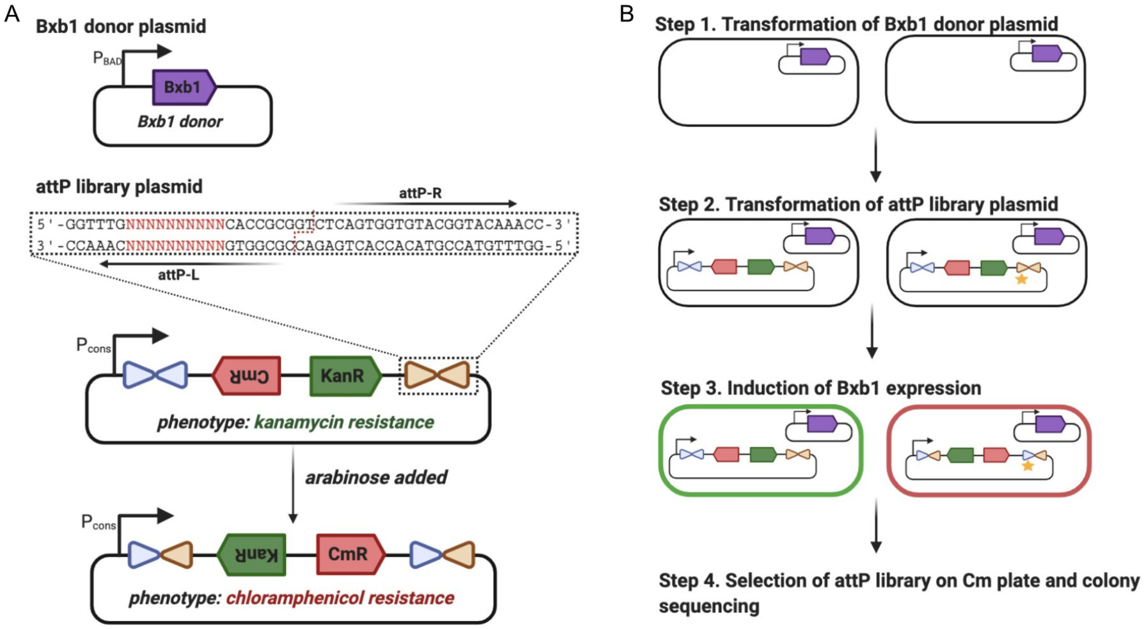


C

| Colony number | Sanger sequencing result |
|---|---|
| 1 | GTCCCCAAAA |
| 2 | GTGTATCGGC |
| 3 | GTTCACGAAA |
| 4 | GTTAGTAAGA |
| 5 | GTAATACTAA |
| 6 | GTTTCCCCGA |
| 7 | GTCGATGACG |
| 8 | GTGCAACCAG |
| 9 | GTTATGCTGA |
| 10 | GTAACCCATA |
| 11 | GTTACCACTA |
| 12 | GTGTGCGACC |
| 13 | ATTAATCAGA |
| 14 | GTCCGTTGTA |
| 15 | GTCGCACACA |
| 16 | GTGGTCCGCA |
| 17 | GCTCAGTCCA |
| 18 | GTGCCGCGAC |
| 19 | GTGCTCTGCG |
| 20 | GTTGACTCAA |

**Figure S1**. *In vivo* selection in *E. coli* with library 1. (A) Diagrams of the Bxb1 donor plasmid (top) and KanR/CmR reporter plasmid (middle). The attP-L half-site library 1 was inserted into the KanR/CmR reporter plasmid for selection. The kanamycin resistance gene and the chloramphenicol resistance gene were placed in opposite directions and downstream of a constitutive promoter. Before addition of L-arabinose to induce Bxb1 recombinase expression, the reporter plasmid confers a kanamycin-resistant phenotype (green). After addition of L-arabinose, a reporter plasmid possessing a functional and efficient variant of the attP-L half-site switches phenotype from kanamycin-resistant (green, middle) to chloramphenicol resistance (red, bottom), and can therefore be selected on LB-agar plates with chloramphenicol. (B) Dual-step transformation of the Bxb1 donor plasmid and the KanR/CmR reporter plasmid, and the phenotype conversion from kanamycin resistance (green) to chloramphenicol resistance (red) after inducible expression of the Bxb1 recombinase. (C) Sanger sequencing results for single colonies selected on chloramphenicol-containing LB-agar plates.

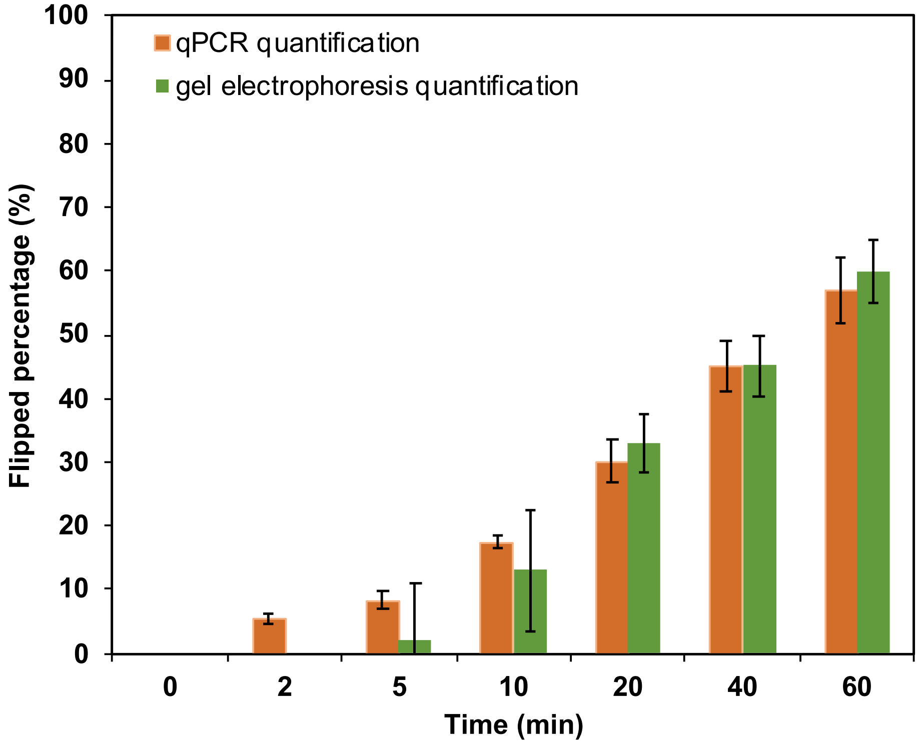


**Figure S2.** Comparison of quantitative methods for measuring DNA inversion. The DNA recombination assay was performed by mixing 20 nM Bxb1 and 10 nM DNA in a 50 µl reaction. The reaction was incubated at 30 °C and terminated by heating after different times. For qPCR quantification, the reaction mixture was diluted and added to a qPCR reaction as a template, then the flipped percentage was calculated based on the $C_q$ value. For gel electrophoresis, HindIII-HF was added to the cooled reaction buffer to digest the DNA. After incubation at 37 °C for 1 h, 15 µl of the digest was subjected to gel electrophoresis. The percentage of flipped DNA was then calculated based on gel quantification of the band intensities of flipped and unflipped DNA. When the DNA flipping reaction duration was 2 min or less, flipped DNA was not detectable by gel quantification. The error bars represent three technical replicates.

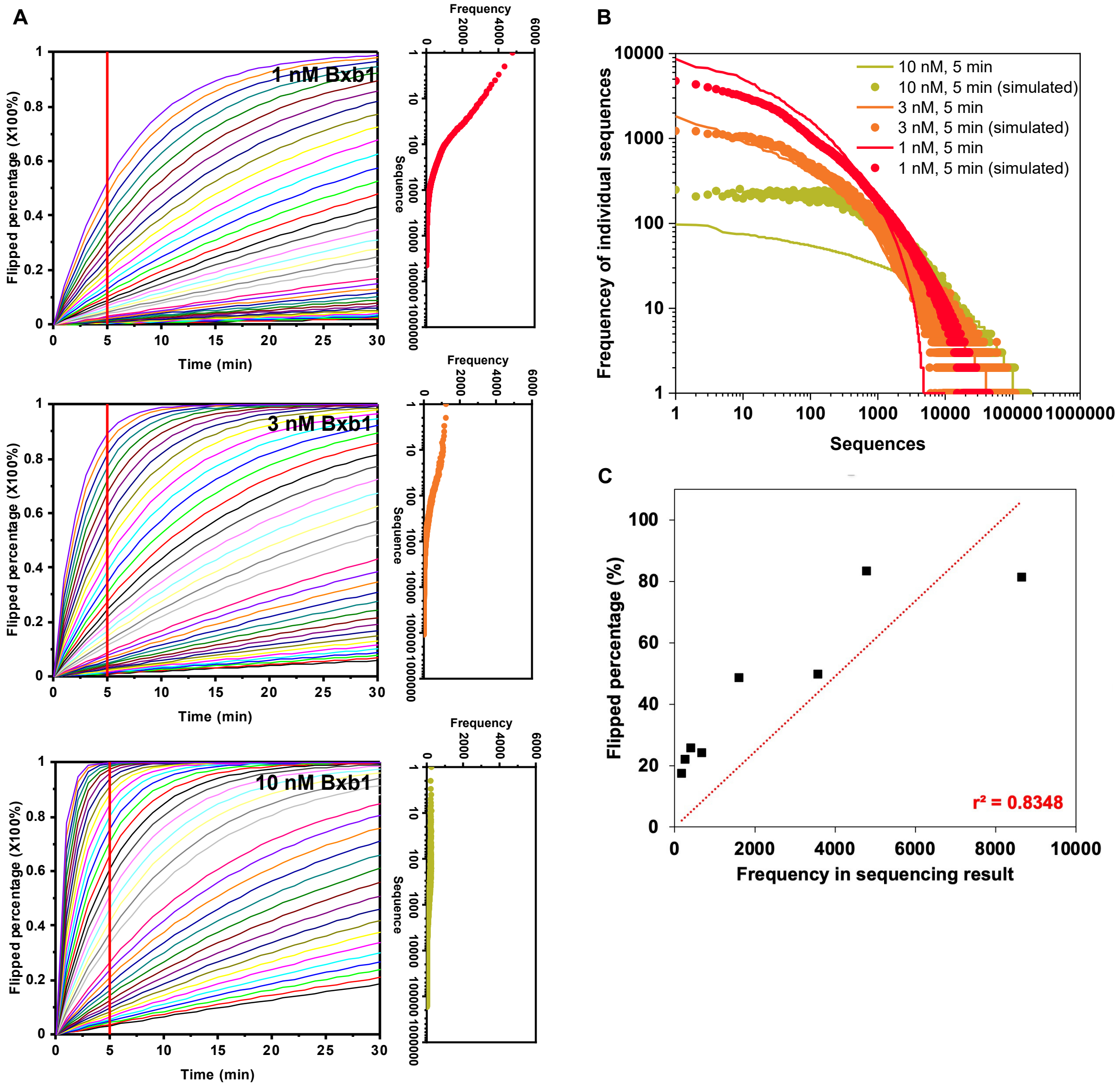


**Figure S3**. Correlation between the frequency in NGS results and the DNA flipping rate constant $k_{flip}$ for a given sequence. (A) (*left*) Simulations of flipped percentage for each possible sequence in library 2 were performed at a total DNA concentration of 10 nM and different Bxb1 concentrations (1 nM, 3 nM, and 10 nM) with first-order reaction kinetics (see Supplementary Method 2), with a representative subset of the response curves shown here. (*right*) At 5 min (corresponding to the vertical red lines in the plots on the left), the frequencies of flipped sequences were computed and plotted as a function of sequence number. (B) Comparison of frequency profiles of the experimental results from NGS with the simulated results based on predicted flipping rate constants $k_{flip}$. (C) Correlation between the flipped percentage in inversion assays conducted on eight individual sequences and the frequency of each of these sequences in the NGS results.

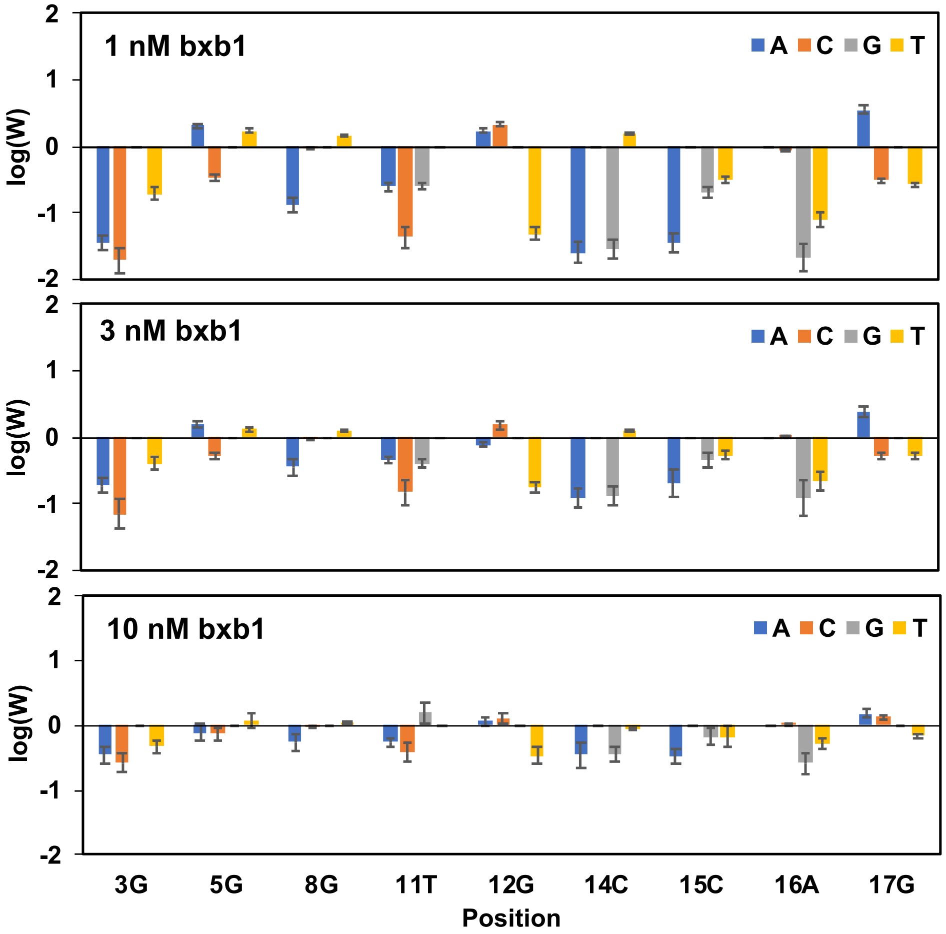


**Figure S4**: Comparison of derived weight scores for different Bxb1 concentrations. Weight score values, W, were generated from three sets of 200 training sequences, randomly chosen from the 3000 most enriched sequences. Error bars represent three randomly chosen 200-sequence training data sets for each Bxb1 concentration.

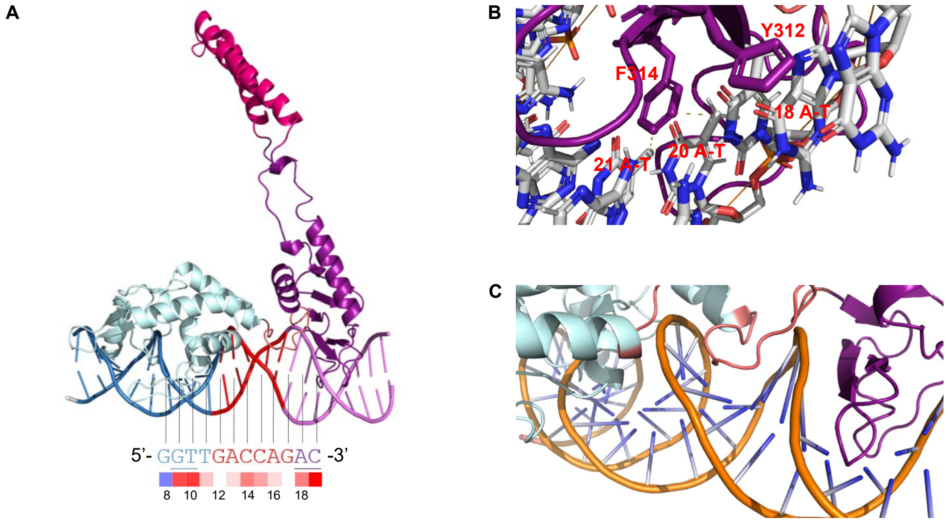


**Figure S5:** Posited interactions between Bxb1 and attP-L based on generated weight scores and homology modeling. (A) The Bxb1-DNA complex structure was simulated using SWISS-MODEL (https://swissmodel.expasy.org/)[1] and NPDock (http://genesilico.pl/NPDock)[2]. (B) Interactions between side chains in the Bxb1 zinc ribbon domain and bound, conserved nucleotides. (C) Interactions between side chains in the Bxb1 linker domain and bound, substitutable nucleotides.

We sought to gain biological insights from the sequence weight scores by comparing our identified conserved positions, positive substitutions, and negative substitutions with previous studies on large serine recombinases.[3] Although a crystal structure of the Bxb1-DNA complex was not available, there was a co-crystal structure of DNA bound by the integrase A118, which is from the same LSR subgroup as Bxb1.[3] Since the C-terminal domain is moderately conserved in LSRs, we used the structure of the A118-attP-L half-site complex (PDB: 4KIS) as a template to simulate Bxb1-DNA interactions with SWISS-MODEL and NPDock. As shown in Fig. S5A, the recombinase domain (RD) and the zinc ribbon domain (ZD) form extensive direct contacts with the 24 bp of the attP-L half-site.

Nucleotide positions 18-24 (purple) at the site terminus are highly conserved among the four half-sites. We therefore hypothesized that these sites are likely essential for function, which was supported by our initial library selections. In addition, a previous study demonstrated that the outside flanking sequences were important for Bxb1 enzyme binding.[4] According to the published report on the A118-attP crystal structure complex,[3] hydrophobic interactions between the side chains of T288(β11), Y295(β12) and Y297(β12) and the 5-methyl groups of 23 A-T, 24 A-T, and 25 A-T are functionally important for protein binding to DNA. It was demonstrated that residues such as tyrosine can interact hydrophobically with the 5-methyl group on thymine, and the carbonyl group of aspartate residues can form hydrogen bonds with the amino group of the nucleic acid. Analogously, in the homology model of Bxb1-attP (Fig. S5B), 18 A-T, 20 A-T, and 21 A-T are suggested to form hydrophobic interactions with Y312 and F314 in the corresponding β-sheets and thus are likely highly conserved (which we experimentally demonstrated for 18A).

This structural model also suggests that nucleotides bound to RD (2-11, blue) and ZD-RD linker (12-16, red) are more tolerant to sequence substitutions. At positions 5, 12, and 17, some substitutions can even enhance the flipping rate. This improvement in activity could arise from higher protein-DNA binding affinity or any following steps in Fig. 1A. The highest weight score, for the G17A substitution, is also supported by a previous study on the Bxb1 attP site in which it was found that the binding affinity of the Bxb1 CTD for the attP-L half-site mutation G17A was twice that of the WT attP-L half-site (70 nM vs. 140 nM).[5] The structural basis of this affinity enhancement is not evident from the homology model.

### Supplemental Methods

Supplemental Method 1: *In vivo* selection using *E. coli*

The low copy number vector p15A-$P_{cons}$-kanR/cmR was used for KanR/CmR reporter plasmid construction and the high copy number pUC19-$P_{BAD}$-bxb1 vector was used for Bxb1 donor plasmid construction by restriction enzyme digestion and ligation. The primers and plasmids used were listed in Table S1 and Table S2.

A two-step transformation method was used to create the selection system. First, the Bxb1 donor plasmid was transformed into TOP10 pro *E. coli* cells. Then, homemade competent cells with the Bxb1 donor plasmid were transformed by the second plasmid p15A-$P_{cons}$-kanR/cmR, which encoded the attP-L library. After the second transformation step, the cells recovered in 950 μL pre-warmed SOB medium and were then incubated at 37 °C and 200 rpm for 1 h. In order to induce Bxb1 protein expression, 0.1% arabinose was added to the culture. After 45 min or 75 min, the cells were washed twice with SOB medium, and then serially diluted before plating on LB-agar plates with antibiotics (50 μg/ml carbenicillin and 15 ug/ml chloramphenicol). The selection plates were incubated at 37 ºC for 14-16 h and colonies that grew on the selection plates were picked for colony PCR using Phusion High-Fidelity polymerase. The amplicons from individual colonies were Sanger sequenced.

Supplemental Method 2: High-throughput reaction rate characterization *in vitro* using NGS

**Species in the *in vitro* DNA library flipping reaction**

| *Species* | *Description* | *Initial concentration* |
|---|---|---|
| S | total DNA concentration of the attP library | 10 nM |
| $S_n$ | DNA concentration of unflipped attP variant n, n=1~1000000 | 0.01 pM |
| $P_n$ | DNA concentration of flipped attP variant n, n=1~1000000 | 0 pM |
| E | Bxb1 protein concentration | 2-300 nM |
| $ES_n$ | Complex of Bxb1 and unflipped attP variant n | 0 pM |

**Parameters in the DNA flipping reaction**

| *Species* | *Description* |
|---|---|
| $k_{cat}(n)$ | catalyst rate constant for attP variant n |
| $k_{on}(n)$ | binding rate constant for attP variant n |
| $k_{off}(n)$ | dissociation rate constant for attP variant n |
| $K_d(n) = k_{off}(n)/k_{on}(n)$ | equilibrium dissociation constant for the Bxb1-attP(n) complex |
| $k_{flip}(n)$ | DNA flipping rate constant for attP variant n |

For attP variant n:

$$S_n + E \underset{k_{off}(n)}{\overset{k_{on}(n)}{\rightleftarrows}} ES_n \xrightarrow{k_{cat}(n)} E + P_n$$

(1) $\frac{dS_n}{dt} = -k_{on}(n) \cdot S_n \cdot E + k_{off}(n) \cdot ES_n$

(2) $\frac{dP_n}{dt} = k_{cat}(n) \cdot ES_n$

(3) $\frac{dES_n}{dt} = k_{on}(n) \cdot S_n \cdot E - k_{off}(n) \cdot ES_n - k_{cat}(n) \cdot ES_n$

(4) $S_n(0) = S_n + ES_n + P_n$

(5) $E(0) = E + \sum ES_n$

In our *in vitro* experiments for quantification of flipping rate, the Bxb1 concentration is 3 nM, the total DNA library concentration is 10 nM, and the wild-type attP-Bxb1 dissociation constant is 60 nM. As shown below, these values allow for two reasonable assumptions: a) quasi-steady state for $ES_n$ and b) $E \approx E(0)$. With these assumptions, the reaction rate can be derived from equation (6) and expressed using equations (7) and (8).

(6) $\frac{dES_n}{dt} = k_{on}(n) \cdot S_n \cdot E(0) - k_{off} \cdot ES_n - k_{cat}(n) \cdot ES_n = 0$

The rate of DNA flipping ($dP_n/dt$) can be written in terms of a DNA flipping rate constant $k_{flip}(n)$:

(7) $\frac{dP_n}{dt} = k_{flip}(n) \cdot E(0) \cdot S_n$

where $k_{flip}(n)$ is a function of $k_{on}(n)$, $k_{off}(n)$, and $k_{cat}(n)$:

(8) $k_{flip}(n) = \frac{k_{cat}(n) \cdot k_{on}(n)}{k_{off}(n) + k_{cat}(n)}$

For the first assumption, the criteria for the validity of the steady-state approximation is that the time scale for the complex ES to reach steady state, $\tau(ES_n)$, should be much shorter than that of S or P, $\tau(S_n)$. This means that, as species concentrations change over the reaction, at every time point, the ES complex can reach equilibrium rapidly. This criterion requires that equation (9) be satisfied[6]:

(9) $\frac{E(0)}{K_m + S} \ll 1$

This equation holds true for our experimental conditions, since the Bxb1 concentration (3 nM) is much less than the wild-type Bxb1-DNA $K_d$ (60 nM). Even for attP-L variants in the DNA library with a several-fold increased affinity for Bxb1, the enzyme concentration would still be much lower than this $K_d$.

Based on the validity of the first assumption, $ES_n$ are at steady state:

(10) $\frac{dES_n}{dt} = k_{on}(n) \cdot S_n \cdot (E(0) - \sum ES_n) - k_{off}(n) \cdot ES_n - k_{cat}(n) \cdot ES_n = 0$

Then, equation (11) can be derived from equation (10):

(11) $\frac{\sum ES_n}{(E(0) - \sum ES_n)} = \sum \left\{ \frac{S_n}{\frac{k_{off}(n) + k_{cat}(n)}{k_{on}(n)}} \right\} < \frac{\sum S_n}{\left[\frac{k_{off}(n) + k_{cat}(n)}{k_{on}(n)}\right]_{min}}$

In our experiments, $\sum S_n$ = 10 nM, which is considerably smaller than the WT attP-Bxb1 $K_d$ = 60 nM, so the second assumption that $E \approx E(0)$ is also reasonable.

Supplemental Method 3: Mathematical simulation of GFP and mCherry co-expression in *E. coli*

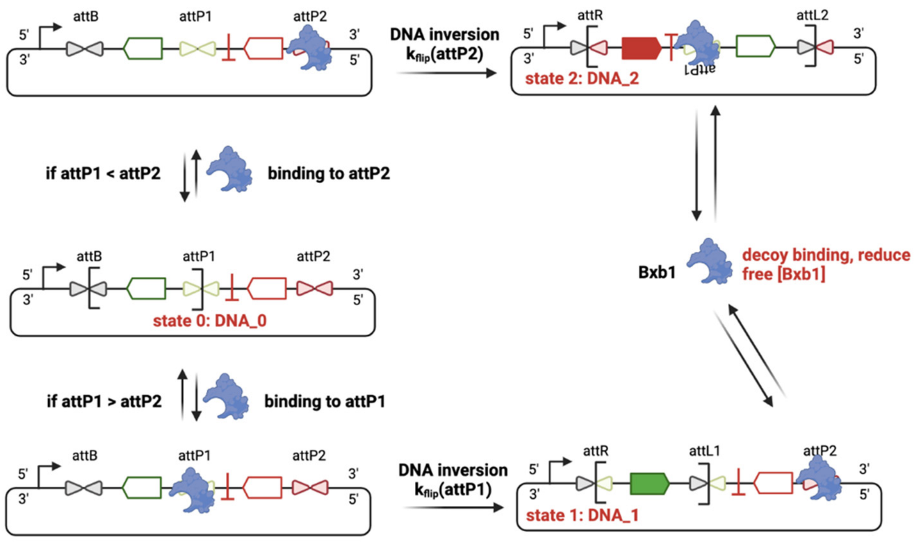


**Species in *E. coli***

| *Species* | *Description* | *Initial concentration* |
|---|---|---|
| x | *E. coli* density | $1.2\times10^9$ cells/ml |
| DNA_Bxb1 | DNA concentration of plasmid DNA coding Bxb1 | 0.25 nM |
| DNA_0 | unflipped reporter plasmid DNA concentration | 25 nM |
| DNA_1 | attP1 flipped reporter plasmid DNA concentration | 0 nM |
| DNA_2 | attP2 flipped reporter plasmid DNA concentration | 0 nM |
| mRNA_Bxb1 | Bxb1 mRNA concentration | 0 nM |
| mRNA_GFP | GFP mRNA concentration | 0 nM |
| mRNA_mCherry | mCherry mRNA concentration | 0 nM |
| Bxb1 | Bxb1 protein concentration | 0 nM |
| GFP | GFP protein concentration | 0 nM |
| mCherry | mCherry protein concentration | 0 nM |

**Parameters in the mathematical model**

| *Parameters* | *Description* | *Values* | *References* |
|---|---|---|---|
| $k_{flip}$(attP1) | DNA flipping rate constant of attP1 | 0.001-0.01 $nM^{-}min^{-1}$ | This study |
| $k_{flip}$(attP2) | DNA flipping rate constant of attP2 | 0.001-0.01 $nM^{-}min^{-1}$ | This study |
| $K_{d.DNA}$(attP1) | Dissociation constant of Bxb1 and attP1 | 10-1000 nM | [5] |
| $K_{d.DNA}$(attP2) | Dissociation constant of Bxb1 and attP2 | 10-1000 nM | [5] |
| $k_m$(Bxb1) | Bxb1 mRNA transcription rate constant | 1.1 $min^{-1}$ | [7] |
| $k_m$(GFP) | GFP mRNA transcription rate constant | 2.2 $min^{-1}$ | [7] |
| $k_m$(mCherry) | mCherry mRNA transcription rate constant | 2.2 $min^{-1}$ | [7] |
| $k_p$(Bxb1) | Bxb1 protein translation rate constant | 0.32 $min^{-1}$ | [8] |
| $k_p$(GFP) | GFP protein translation rate constant | 0.66 $min^{-1}$ | [8] |
| $k_p$(mCherry) | mCherry protein translation rate constant | 0.68 $min^{-1}$ | [8] |
| $k_{deg}$(mRNA) | mRNA degradation rate constant | 0.14 $min^{-1}$ | [7] |
| $k_{deg}$(Bxb1) | Bxb1 protein degradation rate constant | 0.012 $min^{-1}$ | [9] |
| $k_{deg}$(GFP) | GFP protein degradation rate constant | 0.002 $min^{-1}$ | [9] |
| $k_{deg}$(mCherry) | mCherry protein degradation rate constant | 0.002 $min^{-1}$ | [9] |
| μ | *E. coli* growth rate | $3.6\times10^{-3}$ $min^{-1}$ | This study |
| K | maximum cell density | $2.3\times10^9$ cells/ml | This study |

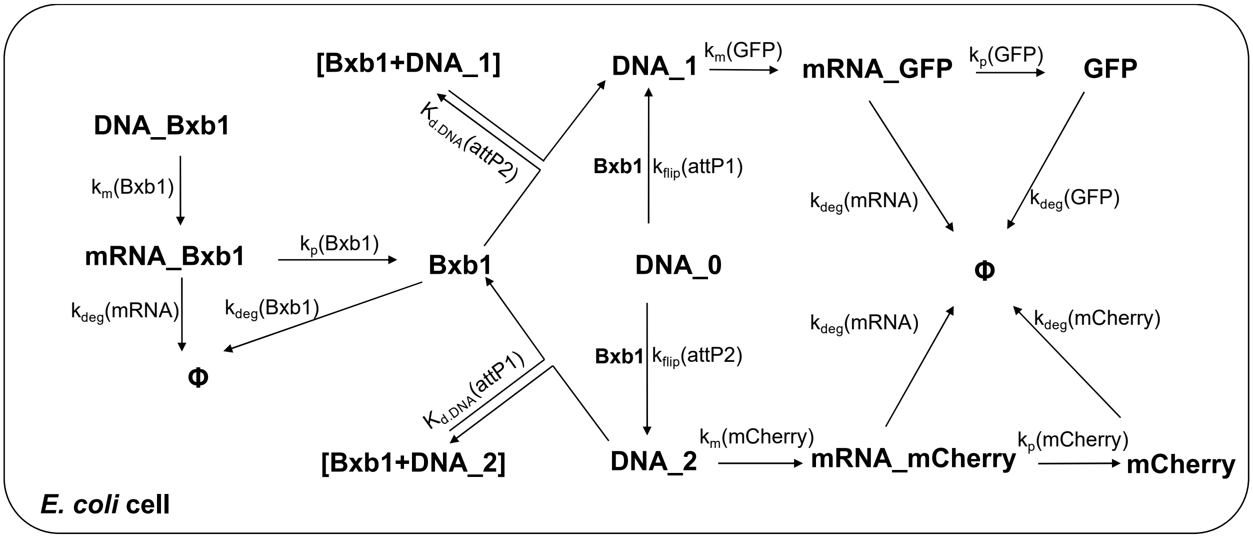


Differential equations describing GFP and mCherry co-expression:

(1) $$\frac{dx}{dt} = \mu \cdot (1 - \frac{x}{K}) \cdot x$$

(2) $$\frac{d(DNA\_Bxb1)}{dt} = -DNA\_Bxb1 \cdot (\mu \cdot (1 - \frac{x}{K}))$$

(3) $$\frac{d(DNA\_1)}{dt} = kflip(attP1) \cdot \frac{Bxb1}{(1 + \frac{DNA\_1}{K_{d\_DNA}(attP2)} + \frac{DNA\_2}{K_{d\_DNA}(attP1)})} \cdot DNA\_0 - DNA\_1 \cdot (\mu \cdot (1 - \frac{x}{K}))$$

(4) $$\frac{d(DNA\_2)}{dt} = kflip(attP2) \cdot \frac{Bxb1}{(1 + \frac{DNA\_1}{K_{d\_DNA}(attP2)} + \frac{DNA\_2}{K_{d\_DNA}(attP1)})} \cdot DNA\_0 - DNA\_2 \cdot (\mu \cdot (1 - \frac{x}{K}))$$

(5) $$\frac{d(DNA\_0)}{dt} = -\frac{d(DNA\_1)}{dt} - \frac{d(DNA\_2)}{dt} - DNA\_0 \cdot (\mu \cdot (1 - \frac{x}{K}))$$

(6) $$\frac{d(mRNA\_Bxb1)}{dt} = k_m(Bxb1) \cdot DNA\_Bxb1 - k_{deg}(mRNA) \cdot mRNA\_Bxb1 - mRNA\_Bxb1 \cdot (\mu \cdot (1 - \frac{x}{K}))$$

(7) $$\frac{d(mRNA\_GFP)}{dt} = k_m(GFP) \cdot DNA\_1 - k_{deg}(mRNA) \cdot mRNA\_GFP - mRNA\_GFP \cdot (\mu \cdot (1 - \frac{x}{K}))$$

(8) $$\frac{d(mRNA\_mCherry)}{dt} = k_m(mCherry) \cdot DNA\_2 - k_{deg}(mRNA) \cdot mRNA\_mCherry - mRNA\_mCherry \cdot (\mu \cdot (1 - \frac{x}{K}))$$

(9) $$\frac{d(Bxb1)}{dt} = k_p(Bxb1) \cdot mRNA\_Bxb1 - k_{deg}(Bxb1) \cdot Bxb1 - Bxb1 \cdot (\mu \cdot (1 - \frac{x}{K}))$$

(10) $$\frac{d(GFP)}{dt} = k_p(GFP) \cdot mRNA\_GFP - k_{deg}(GFP) \cdot GFP - GFP \cdot (\mu \cdot (1 - \frac{x}{K}))$$

(11) $$\frac{d(mCherry)}{dt} = k_p(mCherry) \cdot mRNA\_mCherry - k_{deg}(mCherry) \cdot mCherry - mCherry \cdot (\mu \cdot (1 - \frac{x}{K}))$$

This system of ordinary differential equations was solved in Matlab using ode23s.
